# The evolutionary origin of host association in an ancient bacterial clade

**DOI:** 10.1101/2021.08.31.458344

**Authors:** Max E. Schön, Joran Martijn, Julian Vosseberg, Stephan Köstlbacher, Thijs J. G. Ettema

**Affiliations:** Department of Cell and Molecular Biology, Science for Life Laboratory, Uppsala University; Uppsala, Sweden; Department of Medical Biochemistry and Microbiology, Uppsala University; Uppsala, Sweden; Department of Biochemistry and Molecular Biology, Dalhousie University; Halifax, Canada; Theoretical Biology and Bioinformatics, Department of Biology, Utrecht University; Utrecht, The Netherlands; Laboratory of Microbiology, Wageningen University and Research; Wageningen, The Netherlands

## Abstract

The evolution of obligate host-association of bacterial symbionts and pathogens remains poorly understood. The Rickettsiales represent an order of obligate alphaproteobacterial endosymbionts and parasites that infect a wide variety of eukaryotic hosts, including humans, livestock, insects and protists. Induced by their host-associated lifestyle, Rickettsiales genomes have undergone reductive evolution, leading to small, AT-rich genomes with limited metabolic capacities. We describe several genomes of deep-branching, environmental alphaproteobacteria that branch basal to previously sampled Rickettsiales, and whose genome content are reminiscent of free-living and biofilm-associated lifestyles. Ancestral genome content reconstruction across the Rickettsiales tree revealed that the free-living to host-association transition of this group occurred more recently than previously anticipated, and likely involved the repurposing of a type IV secretion system.

**One-Sentence Summary:** Deep-branching Rickettsiales provide insights into the evolution of obligate host-associated lifestyle

## Main Text

Bacterial pathogens represent a leading cause of human, livestock and crop disease, leading to significant economic loss worldwide. While molecular and cellular mechanisms of microbial pathogenesis have been described into considerable detail, the origin and emergence of bacterial pathogenicity remain poorly understood. The Rickettsiales represent a widespread and diverse order of obligate host-associated alphaproteobacteria that infect a wide variety of eukaryotic species, including protists, leeches, cnidarians, arthropods, and mammals (*1*). Well-known examples include *Rickettsia prowazekii*, the causative agent of epidemic typhus in humans (*2*), and *Wolbachia*, a genus of bacteria infecting over two-thirds of all arthropods and nearly all filarial nematodes (*3*). Rickettsiales genomes have been shaped by ongoing reductive evolution, driven by their host’s nutrient-rich cytoplasm, enhanced genetic drift because of small effective population sizes and frequent bottlenecks, and an uninhibited Muller’s ratchet (*4*). As a result, their genomes are typically small (<1.5 Mb), AT-rich (<40% GC), display a low coding density (<85%), lack metabolite biosynthesis genes and display a high degree of pseudogenization (*5*). Rickettsiales employ various host-interaction factors, including a type IV secretion system (T4SS), various associated host cell manipulating effector proteins (*1, 6, 7*) and an ATP/ADP translocase, which facilitates energy parasitism by exchanging host cell ATP for endogenous ADP (*8, 9*). Beyond this, the lineages belonging to the currently recognized Rickettsiales families (Rickettsiaceae, Anaplasmataceae, Midichloriaceae and Deianiraeaceae) each have adopted specific lifestyles to interact with their respective host cell environment (*1, 10–13*). It is, however, unclear how and when the Rickettsiales ancestor transitioned from a free-living to a host-associated lifestyle.

To shed light on the emergence and early evolution of Rickettsiales, including their host associated lifestyles, we screened publicly available metagenomic repositories for the existence of deep-branching rickettsial genomes (see SOM for details, Data S1, Data S2). This analysis, which was based on the phylogenetic screening of metagenomic contigs containing at least five genes of a conserved 15-ribosomal protein gene cluster (see SOM for details), resulted in the reconstruction and identification of several putative Rickettsiales metagenome-assembled genomes (MAGs) derived from aquatic (marine, lake, aquifer and tailings water) environments (Fig. S1). To assess the phylogenetic affiliation of these MAGs in more detail, and to exclude the possibility that these were derived from mitochondrial sequences, we performed a phylogenomic analysis based on a previously established set of 24 mitochondrially encoded marker genes (*14*). We found that the identified lineages represented three distinct, novel Rickettsiales clades that were unrelated to mitochondria (Fig. 1, Fig. S2). In addition, the analysis provides further support for a mitochondrial origin outside of the currently sampled alphaproteobacterial diversity (*14, 15*).

**Fig. 1.**
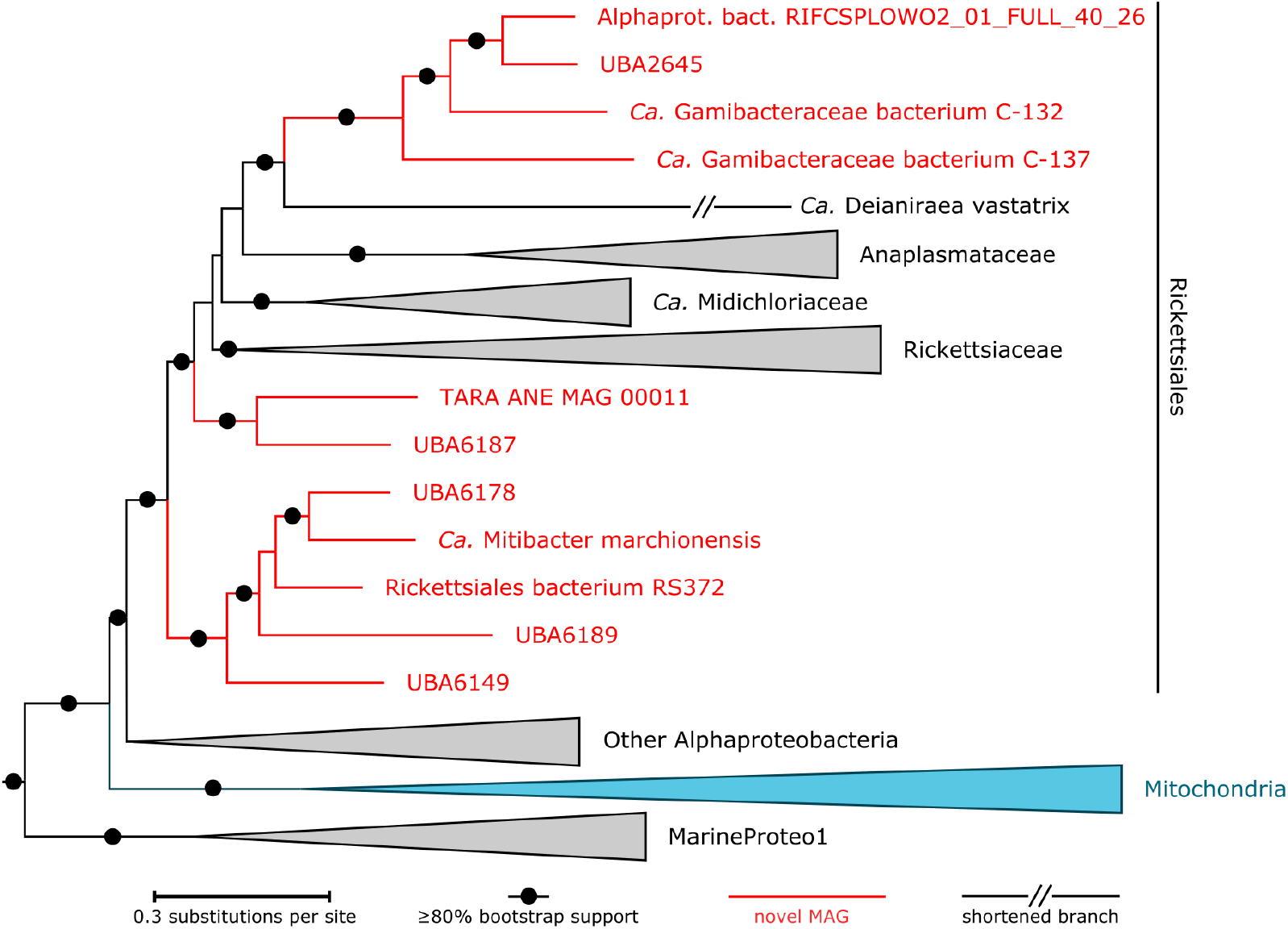
Identification and phylogenomics of novel Rickettsiales-associated alphaproteobacteria. Phylogenetic tree based on the ‘alphamito24’ dataset including novel Rickettsiales MAGs and other recently sequenced genomes (see SOM for details). The 20% most heterogeneous sites were removed before tree inference with IQTREE under the PMSF approximation LG+C60+F+Γ4 model (guidetree: LG+F+Γ4) and 100 non-parametric bootstraps. Tree was rooted with Beta-, Gammaproteobacteria and Magnetococcales. See Fig. S2 for the uncollapsed tree.

To pinpoint the phylogenetic position of the novel Rickettsiales-associated clades, we performed a more in-depth phylogenomic analysis based on a dataset comprising 116 marker proteins (*14*) (Fig. S3; see SOM for details). Intriguingly, this analysis confirmed that two of the three clades (hereafter Mitibacteraceae and Athabascaceae; see SOM for taxonomic descriptions), were found to branch as sister groups to all currently recognized Rickettsiales families, with Mitibacteraceae representing the deepest branching clade (Fig. S4, Fig. S5, Data S1). These phylogenetic placements were robust to removal of the most heterogeneous sites (Fig. 2A, Fig. S6, Fig. S7, Tab. S1; see SOM for details). The third Rickettsiales-associated clade (hereafter Gamibacteraceae; see SOM for taxonomic description) formed a distinct sister group of ‘*Candidatus* Deianiraea vastatrix’ (Fig. 2A; see SOM for details), a recently described Rickettsiales lineage displaying a unique host-associated extracellular lifestyle, including the ability to replicate outside *Paramecium* host cells (*11*). An analysis of 16S rRNA gene sequences found in Gamibacteraceae and Mitibacteraceae MAGs (none were identified in Athabascaceae MAGs) revealed an environmental distribution exclusive to aquatic habitats. While close homologs of the *Ca*. Mitibacter marchionensis 16S rRNA gene were only identified in marine environments (Fig. 2B; Fig. S8), Gamibacteraceae-related sequences were also observed in lakes and aquifers (Fig. 2C; see SOM for details).

**Fig. 2.**
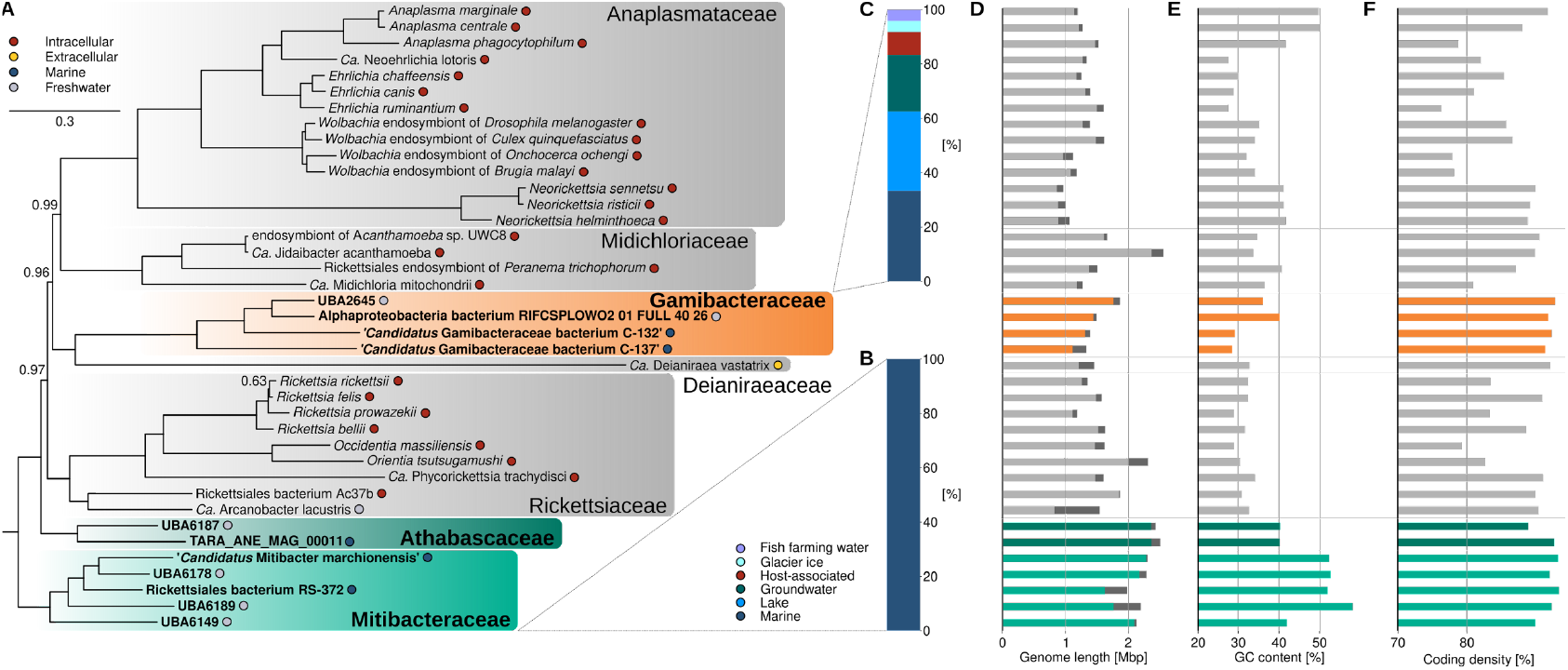
Phylogeny and comparative analysis of deep-branching Rickettsiales genomes. (A) Rickettsiales species tree based on a dataset of 116 marker genes highlighting the phylogenetic position of Gamibacteraceae (orange shading), Athabascaceae (dark green shading) and Mitibacteraceae (light green shading). The tree was inferred using PhyloBayes (CAT+LG+Γ4, 20000 cycles with a burn-in of 5000 cycles, see SOM for details). Support values correspond to posterior probabilities and are only shown for values below 1. See Fig. S6 for the complete tree including the outgroup. Distribution of environmental Mitibacteraceae (B) and Gamibacteraceae sequences (C) obtained from the NCBI nt database (see SOM for details). Colors represent the sequences’ source environments. For each genome the observed genome size in Mbp (D), with estimated genome size in dark grey shading, the GC-content (E) and the coding density, calculated as the total length of protein-coding genes divided by the genome size (F), are shown.

Compared to obligate host-associated Rickettsiales (hereafter referred to as ‘classical Rickettsiales’; including Gamibacteraceae), the reconstructed genomes of Mitibacteraceae and Athabascaceae are, on average, larger in size, display a higher GC content and a higher protein coding density (Fig. 2D-F, Data S3). In addition, they uncharacteristically encode a full biosynthetic capacity for all amino acids, purines and pyrimidines, and nearly all subunits of the main oxidative phosphorylation complexes. Like classical Rickettsiales, they encode a complete tricarboxylic acid cycle but lack a complete glycolysis pathway. However, they do encode a complete glyoxylate cycle, providing the genetic potential to use substrates like acetate as sole carbon source (Fig. S9). Mitibacteraceae and Athabascaceae genomes do not, in contrast to other aquatic alphaproteobacteria, encode proteorhodopsin-related genes, indicating that they lack the capacity to generate energy from a light-driven H^+^ gradient. We furthermore found that Mitibacteraceae and Athabascaceae encode a flagellum and associated chemotaxis machinery (Fig. 3A), suggesting a possible motile lifestyle. We could not identify genes potentially reflective of an intracellular or parasitic lifestyle, besides those genes encoding a typical Rickettsiales (*rvh*-type) T4SS (Fig. 3BC, Fig. S10). This T4SS is typically used for the manipulation of the host cell via translocation of effector proteins (*6, 7*). We failed to detect proteins such as an ATP/ADP translocase, or an accumulation of proteins with eukaryotic-like domains such as ankyrin repeat (ANK) and leucine rich repeat (LRR) domains that could act as potential effector proteins for the identified T4SS (*12, 16*) (Fig. S11). Hence, while representing deep-branching Rickettsiales relatives, the Mitibacteraceae and Athabascaceae seemingly lack signatures indicative of an obligate host-associated lifestyle typical of classical Rickettsiales. Instead, their genome structure and metabolic capacity are reminiscent of copiotrophic, free-living bacterioplankton (*17, 18*) (Fig. 3A, Fig. 2D-F and Fig. S9).

**Figure 3.**
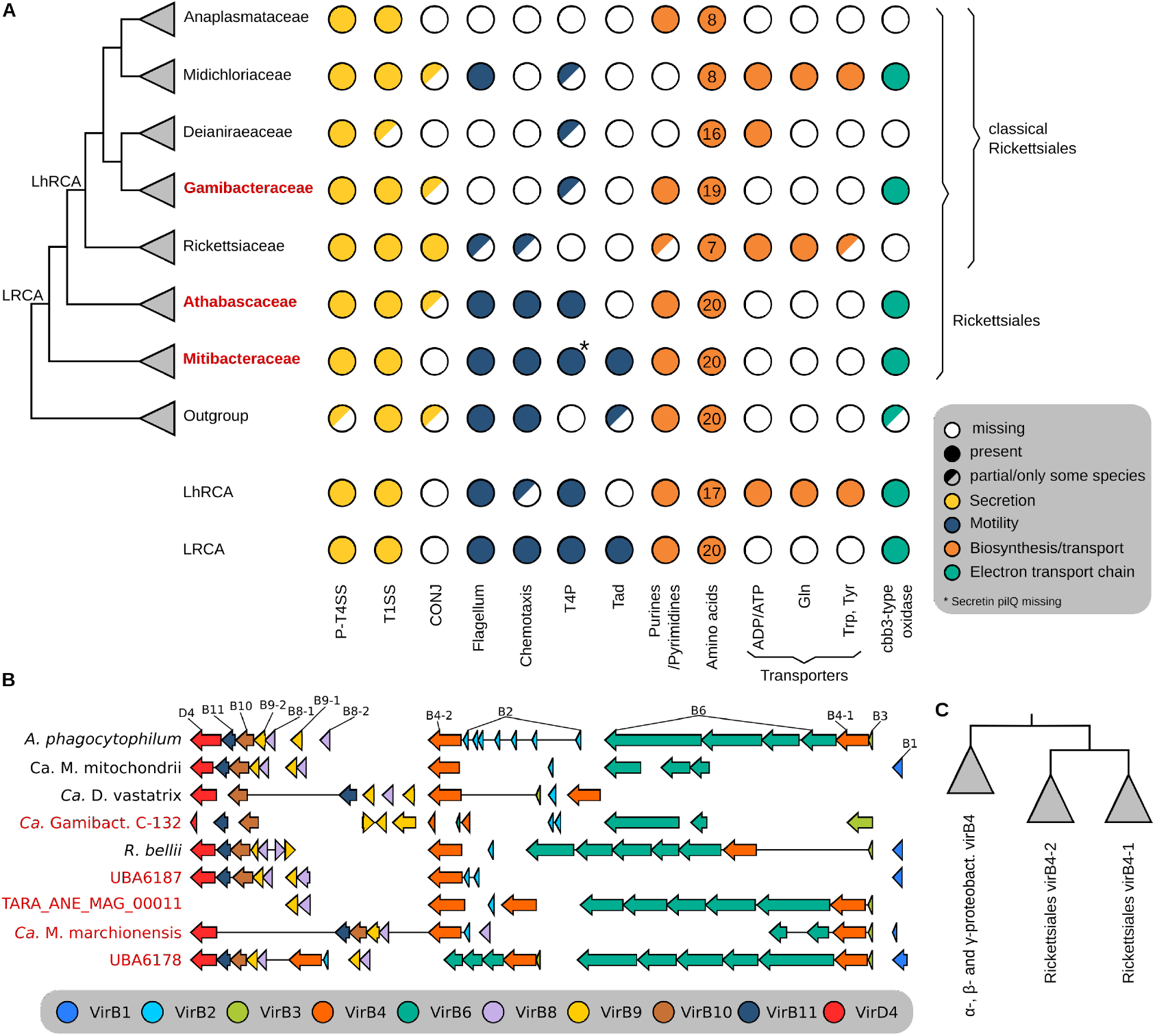
Distribution and ancestral inference of phenotypic traits and pathways across Rickettsiales. (A) Presence and absence of pathways are shown for the seven families of Rickettsiales (including the newly described Gamibacteraceae, Athabascaceae and Mitibacteraceae), the alphaproteobacterial outgroup as well as the last common ancestors of the classical, host-associated Rickettsiales (LhRCA) and of all Rickettsiales (LRCA). Pathways are grouped into ‘secretion’ (*yellow*), ‘motility’ (*blue*), ‘biosynthesis and transport’ (*orange*) and ‘electron transport chain’ (green). In some cases, pathways are either partial (one or several genes missing) or are absent from some species, but present in others. The Gln transporter could potentially also transport other polar amino acids such as Ser, Thr and Asn. (B) Synteny of the rickettsiales-type *vir* homolog (*rvh* T4SS) in classical (black) and environmental (red) Rickettsiales. Taxa are ordered according to the species tree in (A). One representative per family except Athabascaceae and Mitibacteraceae with two representatives each. (C) Schematic phylogenetic tree showing the common duplication of the *vir*B4 gene in LRCA. See Fig. S10 for the uncollapsed tree.

The identification of deep-branching Rickettsiales clades allowed us to study the ancient evolutionary transition from a free-living to a host-associated ancestor in unprecedented detail. We used a gene tree-species tree reconciliation method (*19, 20*) to reconstruct key stages of this transition (see SOM for details) by inferring phylogenetic trees from 4,240 alphaproteobacterial and rickettsial protein families and reconciling these with the previously obtained species tree (Fig. 2A). We then inferred gene duplication, transfer, loss and origination events for all species tree nodes. This allowed us to infer the presence or absence of gene families in these nodes, as well as the relative timing of such events (see SOM and Data S4). We focused on two nodes key to the transition at hand: the last <u>R</u>ickettsiales <u>c</u>ommon <u>a</u>ncestor (LRCA) and the last <u>h</u>ost-associated (classical) <u>R</u>ickettsiales <u>c</u>ommon <u>a</u>ncestor (LhRCA). Ancestors of the different lineages/families within Rickettsiales (Data S6; Fig. S12; see SOM for details) were considered as well. We inferred 1,432 protein coding genes in LRCA, reconstructing a considerably larger ancestral genome than most classical Rickettsiales (Data S5). This is most likely an underestimation due to missing data, as gene families that were present in an ancestor could have gone extinct or have remained unsampled in the genomes analyzed in the present study. Analysis of the inferred ancestral genome content revealed that, like classical Rickettsiales, LRCA displayed typical features of a heterotrophic metabolism and adaptations to aerobic conditions, i.e. oxidative phosphorylation and defense against reactive oxygen species (Fig. S9, Data S6). However, we inferred the potential to *de novo* synthesize all amino acids as well as nucleotides, lacking an ATP/ADP translocase and several amino acid transporters conserved in most classical Rickettsiales and LhRCA (Fig. 3A). While a facultative eukaryote association cannot be fully excluded, several additional lines of evidence support a free-living lifestyle for LRCA. First, LRCA is inferred to have had the potential for sulfate uptake, assimilatory sulfate reduction (Fig. 4; Fig. S9), ammonium uptake and arsenic resistance - all characteristic of typical free-living marine bacterioplankton (*21*). Second, besides a previously proposed ancestral flagellum (*11, 12*), including the necessary machinery for chemotaxis (both of which were retained in LhRCA) (Fig. 3A), we also infer a complete gene set for the biosynthesis of type IV pili (T4P), which, apart from motility, can function in surface colonization as a first step towards biofilm formation (*22*), and in evasion of predation by protist grazers (*23*). Finally, we inferred a tight adherence (*tad*) pilus as well as genes related to the *pel* gene cluster for the synthesis of extracellular polysaccharides (*24, 25*) (Fig. 4; Fig. S13AB), both traits of biofilm formation (*26*) that were subsequently lost in LhRCA.

**Figure 4.**
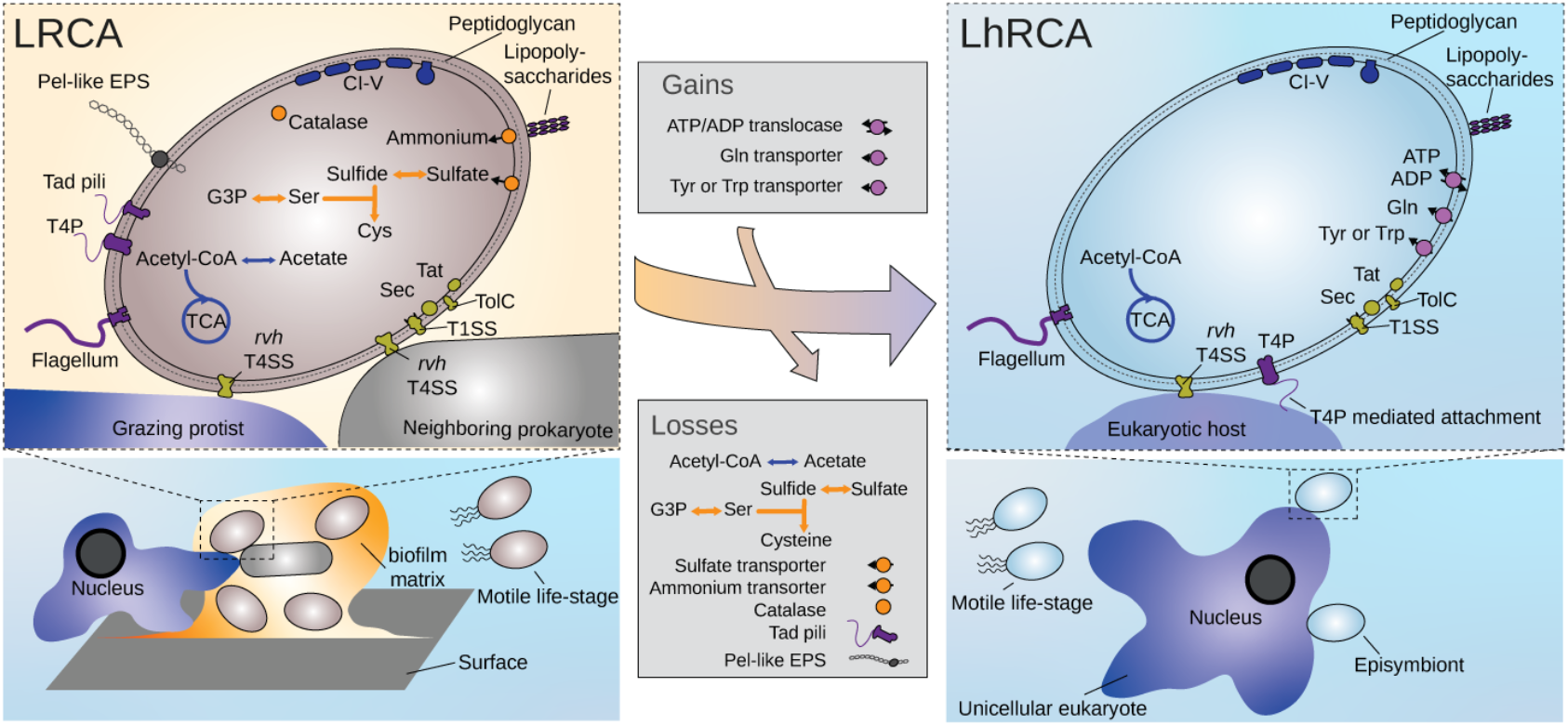
Evolutionary transition from a free-living to a host-associated lifestyle in Rickettsiales ancestors. Proposed lifestyle of the last common ancestors of all Rickettsiales (LRCA) and of the classical, host associated Rickettsiales (LhRCA) in blue boxes at the bottom indicating marine environments. Rickettsiales cells are depicted as ellipsoid cell models in brown and blue for LRCA and LhRCA respectively. Reconstructions of key genomic features of ancestors are depicted in cell models in the respective boxes (see SOM for details). Arrow in the center indicates evolutionary transition from LRCA to LhRCA and grey boxes above and below show gains and losses of genomic traits, respectively. Orange and blue arrows indicate metabolic pathways related to sulfate assimilation or central carbon metabolism, respectively. Genes related to tight adherence (Tad) pili; type 4 pilus (T4P); type I (T1SS) or *rvh* type IV (T4SS) secretion systems; Pel-like exopolysaccharide (EPS) synthesis; tricarboxylic acid (TCA) cycle; electron transport chain complexes (C) I-V.

Based on the considerable metabolic capacity atypical for obligate symbionts, as well as hallmark features of chemotactic motility and biofilm formation, we propose a free-living lifestyle associated with biofilms for LRCA (Fig. 4). The reconstructed genome content is reminiscent of the biphasic life cycle commonly observed in free-living prokaryotes, consisting of a motile planktonic and a surface-attached life stage within biofilms, respectively (*27*) (Fig. 4). While the *rvh*-type T4SS was likely acquired by LRCA through horizontal gene transfer (HGT) from a proteobacterial donor (Fig. 3C, Fig. S10), and subsequently evolutionary conserved throughout Rickettsiales evolution (*6, 7*), we could not identify known T4SS effectors related to eukaryotic host-association in LRCA. We however detected an enrichment of proteins containing peptidoglycan-binding (PGB) domains in the deep branching Rickettsiales genomes (Fig. S11), which could potentially serve as effectors of a ‘bacteria killing’-type T4SS known from other Proteobacteria (*28, 29*), and in analogy with interbacterial competition mediation known from T4SSs (*30*). Yet, PGB domain proteins are also known to operate in other cellular processes, including peptidoglycan synthesis and cell shape remodeling (*31*). Given that cell shape plasticity represents a known mechanism to evade predation by free-living protists (*32*), the observed PGB domain proteins could play a role in this process. Similarly, the T4SS could also function as a defense mechanism against predation by free-living protists, using yet unknown effectors. The latter scenarios would be compatible with the inferred biofilm-forming capacity of LRCA, as aquatic prokaryotes have been shown to form biofilms to evade protist grazing (*32, 33*). Altogether, our results support a scenario in which the repurposing of the T4SS initially used to mediate interbacterial competition or combat protist predation as a plausible intermediate stage towards the inferred obligate host-associated and parasitic lifestyle of LhRCA. While we cannot infer the exact nature of the host interaction of LhRCA, the emergence of the intracellular lifestyle in Rickettsiales was likely preceded by a stage of exo-parasitism conserved in *Ca*. Deianiraea vastatrix (*11*). Here, LhRCA attachment to protist host cells could have been facilitated by the retained T4P, as demonstrated for the predatory bacterium *Vampirococcus lugosii* (*34*). Furthermore, the potential for motility, conserved in extant members of the Midichloriaceae and Rickettsiaceae (*10, 35*) likely functioned in extracellular dispersal between host cells, while the majority of genes related to biofilm formation were lost (Fig. 3A; Fig. 4). The obligate host association of LhRCA was accompanied by extensive genome reduction (Fig. 4, see SOM for details), further exacerbated by extensive genetic drift and limited HGT bias upon adoption of obligate intracellular lifestyles characteristic of most Rickettsiales families.

Recent advances in genome-sequencing technologies and metagenomics approaches have significantly expanded our current views on microbial diversity at the genomic level (*36*). By providing genomic blueprints of a vastness of novel and known bacterial and archaeal clades, studies to reconstruct the evolution of physiological features or lifestyles have now become feasible. Our study reports the discovery of Mitibacteraceae and Athabascaceae, two deep-branching Rickettsiales clades that, based on their genome content, seem to represent free-living relatives of otherwise obligate host-associated Rickettsiales. Subsequent reconstruction of ancestral genome content across the Rickettsiales tree allowed us to reconstruct characteristics of the last common ancestor of Rickettsiales, and identify key stages of the free-living to obligate host association observed in classical parasitic and endosymbiont Rickettsiales. Further studies of the Mitibacteraceae and Athabascaceae, preferably by means of cultivation (*37*), are likely to provide further insights of the physiology and lifestyle of these lineages, and potentially of the last common ancestor of Rickettsiales.

## Supporting information

DataS1

DataS2

DataS3

DataS4

DataS5

## Acknowledgments

We are grateful to those involved in the generation of metagenomic datasets analyzed in the present work and for making these publicly available to the scientific community. We thank the Uppsala Multidisciplinary Center for Advanced Computational Science (UPPMAX) at Uppsala University and the Swedish National Infrastructure for Computing (SNIC) at the PDC Center for High-Performance Computing for providing computational resources.

## Author contributions

Conceptualization: TJGE

Methodology: MES, JM, SK, TJGE

Formal analyses: MES, JM, JV, SK

Investigation: MES, JM, JV, SK

Funding acquisition: JM, TJGE

Visualization: MES, JM, SK

Validation: MES, JM

Supervision: TJGE

Writing – original draft: MES, JM, SK, TJGE

Writing – review & editing: JM, MES, JV, SK, TJGE

## Data and materials availability

In addition to data available in the supplementary materials, files containing sequence datasets, alignments, and phylogenetic trees in Newick format are archived at the digital repository Figshare: <u>10.6084/m9.figshare.c.5494977</u>. MAGs generated in this study are linked to BioProject <u>PRJNA746308</u>. Accessions for genomes analyzed in this study can be found in Data S3. Custom scripts used are available on github (https://github.com/maxemil/rickettsiales-evolution, https://github.com/maxemil/ALE-pipeline and https://github.com/novigit/broCode).

## Supplementary Materials for

### Materials and Methods

#### Sample selection

All publically available Tara Oceans assemblies (*38*) were screened with the RP15 pipeline described by Zaremba-Niedzwiedzka et al. (*39*) for presence of Rickettsiales-related lineages, as described by Martijn et al. (*14*). The RP15 pipeline approximates the phylogenetic position of all taxa present in a metagenome assembly for which at least five out of fifteen well conserved ribosomal proteins are encoded on a single contig. In the end, two assemblies (125_MIX_0.22-3 and 067_SRF_0.22-0.45) were identified (Data S2).

#### Raw sequence data

The raw sequence data from all samples corresponding to the two selected assemblies and from additional samples (Data S2) were downloaded from the Tara Oceans project ERP001736 on the EBI Metagenomics portal.

#### Read preprocessing

All reads were preprocessed similar to Martijn et al. (*14*). SEQPREP was used to merge overlapping read pairs into single reads and remove read-through Illumina adapters, and TRIMMOMATIC v0.35 (*40*) was used to remove residual Illumina adapters, trim low quality base-calls at starts and ends of reads, remove short reads and finally remove reads that had a low average phred score. The overall quality and presence of adapter sequences of processed and unprocessed reads was assessed with FASTQC v0.11.4 (31).

#### Metagenome assembly

The preprocessed metagenomic reads from the two selected samples (125_MIX_0.22-3 and 067_SRF_0.22-0.45) were re-assembled with metaSPAdes (*41*), a mode of SPAdes 3.7.0 with k-mers 21,33,55,77. In case a sample was associated with multiple sequencing runs, all preprocessed reads from the different sequencing runs were pooled prior to assembly.

#### Phylogenetic diversity in metagenome assemblies and public MAG datasets

The RP15 pipeline was used to estimate the phylogenetic diversity of alphaproteobacterial lineages present in the two re-assembled metagenomes and published MAGs of Parks et al. (*42*) (PRJNA348753), Tully et al. (*43*) (PRJNA391943), Delmont et al. (*44*) (https://doi.org/10.6084/m9.figshare.4902923) and Anantharaman et al. (*45*) (PRJNA288027). Only the MAGs categorized by the aforementioned studies as ‘Alphaproteobacteria’ or ‘Rickettsiales’ were considered. Protein coding sequences were predicted with Prodigal v2.60 (*46*). The RP15 pipeline was first run with a reference set of 90 representative bacteria and archaea to identify alphaproteobacterial contigs (Data S1) (*14*). The ribosomal proteins encoded on these contigs were then incorporated in a second RP15 dataset consisting of their orthologs in 84 representative alphaproteobacteria, 12 mitochondria, 2 MarineProteo1, 2 magnetococcales, 4 betaproteobacteria and 4 gammaproteobacteria. A concatenated supermatrix alignment was prepared (alignment: MAFFT-L-INS-i, alignment trimming: TRIMAL -gt 0.5). To reduce compositional bias—a phylogenetic artefact for which alphaproteobacteria are particularly sensitive (*14, 47–49*)—we removed 20% of the sites that contributed most to compositional heterogeneity (*48*). A phylogenetic tree was inferred from the compositionally trimmed supermatrix alignment with IQTREE v1.6.9 (-m LG+C60+F+G -bb 1000 -nm 250; Figure S1) (*50*).

#### Binning of metagenomic contigs

For each metagenome assembly, contigs larger than 2kb were grouped into bins based on differential coverage across samples, tetranucleotide frequency profiles, GC-composition and read-pair linkage as described by (*14*). Briefly, the contigs were cut every 10kb, unless the remaining fragment was shorter than 20kb. Then the preprocessed reads of a set of sequencing runs (125_MIX_0.22-3: all sequencing runs listed in Data S2, 067_SRF_0.22-0.45: sequencing runs ERR598994, -599144, -594313, -594325, -594395 and -594404, Data S2) were mapped onto the fragmented contigs with KALLISTO v0.42.5 (*51*), yielding differential coverage profiles per fragmented contig. This was then used together with tetranucleotide frequency information by CONCOCT v0.4.0 (*52*) to group the fragmented contigs into bins. Bins containing the Rickettsiales ribocontigs (see Fig. 1) were then assessed and cleaned with MMGENOME (*53*) using differential coverage, GC-composition, read-pair linkage and presence of 139 genes well-conserved across Bacteria. Finally, the fragmented contigs of the cleaned bins were replaced by their corresponding full length contigs. In case not all fragmented contigs from a corresponding full length contig were present in a cleaned bin, the full length contig would only be included in the final bin if the majority of the fragmented contigs were present. This yielded three draft genome bins: “BIN125”, “BIN67-1” and “BIN67-3”.

We aimed to improve their quality by recruiting reads from all Tara Oceans metagenomes that had coverage for the draft bins (BIN125: ERR594323, -599156, -594338, -594339, -59434, BIN67-1 & BIN67-3: ERR594395, -594404, -598994, -599144, -594313, -594325) and performing a second round of assembly and binning. This was completed as follows. Preprocessed reads putatively derived from the genomes of interest were recruited from the selected metagenomes by classifying them with Bowtie2 (*54*), and CLARK-S v1.2.3 (*55*) using a set of reference Rickettsiales genomes. The recruited reads were combined in two separate pools, one for BIN125 samples and one for BIN67-1/BIN67-3 samples. Each pool was assembled separately with SPAdes (--careful) (*56*). The final BIN125 bin was obtained by removing all contigs < 3300 bp. The final BIN67-1 and BIN67-3 bins were obtained by separating the contigs ≥ 1500 bp into two groups with CONCOCT (--clusters 4).

#### Completeness and redundancy estimates

We used the miComplete tool v.1.1.1 (*57*) to estimate the completeness and redundancy of the MAGs as well as the reference genomes using a set of bacterial marker genes.

#### Annotation

All bins were annotated with prokka v1.12 (*58*), which was altered to allow for partial gene predictions on contig-edges (GitHub pull request #219), with the options --compliant, -- partialgenes, --cdsrnaolap and --evalue 1e-10, and with barrnap as the rRNA predictor. We used the eggNOG-mapper v1.0.3 to get annotations from the EggNOG database 4.5.1 (from which we gathered the alphaNOGs) (*59, 60*). We assigned KEGG orthology (KO) and enzyme commision (EC) numbers using GhostKOALA (Minoru Kanehisa, Sato, and Morishima 2016). Additionally, we annotated the proteins using the Carbohydrate-Active enZYmes Database (CAZY, using HMMER) (*61, 62*), the Transporter Classification Database (TCDB, using BlastP) (*63, 64*) and used InterProScan to annotate the proteins with PFAM, TIGRFAM and IPR domains (*65–67*). For detailed annotation of secretion systems and filamentous structures we screened proteomes using MacSyFinder (*68, 69*) v2.0rc1 with the “TXSScan” (*68, 69*) and “TFF-SF” (*70*) HMM models with “—db_type unordered”. Finally, we used DIAMOND (*71*) to perform similarity searches of the proteins against NR and recorded the taxonomic annotation of the last common ancestor of all hits within 2 percent of the best score. Further searches of PFAM domain profiles (*67*) were performed using HMMER (*62*) for eukaryotic domains commonly found in Rickettsiales T4SS effectors (*7, 12*) as well as in Xanthomonas bacterial-killing T4SS effectors (*28, 29*). All annotations are presented in Data S6.

#### Alphaproteobacteria and mitochondria species tree

A phylogenomics dataset was constructed by updating the ‘24 alphamitoCOGs with more diverse mitochondria’ dataset from Martijn et al. (*14*) (from here on ‘alphamito24’) with the three Rickettsiales MAGs reconstructed in this study, eight Rickettsiales MAGs identified from public MAG datasets (*42–45*), *Candidatus* Deianiraea vastatrix (*11*), endosymbiont of *Peranema trichophorum* and endosymbiont of *Stachyamoeba lipophora* (*72*) and *Candidatus* Phycorickettsia trachydisci (*73*). Orthologs were identified by PSI-BLAST (*64*) by using the gene alignments of the alphamito24 dataset as queries. Non-orthologs were detected and removed via single gene tree inspections (alignment: MAFFT L-INS-i v7.471 (*74*), alignment trimming: trimAl v1.4.rev15 - gappyout (*75*), phylogenetic inference: IQTREE v1.6.12 -fast -m LG+F+G (*50*)). A supermatrix alignment was prepared from the updated orthologous groups by re-aligning with MAFFT L-INS-i and trimming the alignments with BMGE -m BLOSUM30 (65) prior to concatenation. The alignment was finalized by removing the 20% most compositional heterogeneous sites with the χ^2^-trimmer (*14, 48*). A phylogenetic tree was inferred under the PMSF approximation (*76*) of the LG+C60+F+Γ4 model (guidetree: LG+G+F, 100 non-parametric bootstraps).

#### Rickettsiales species tree

The ‘129-panorthologs’ phylogenomics dataset of Martijn et al. (*35*) was updated with the same MAGs and genomes that were used to update the alphamito24 dataset, as well as the recently sequenced Rickettsiales “UBA6177” (*42*), *Candidatus* Xenolissoclinum pacificiensis (*77*), *Candidatus* Fokinia solitaria (*78*), *Candidatus* Neoehrlichia lotoris (ASM96479v1), *Neorickettsia helminthoeca* (ASM63298v1), *Candidatus* Jidaibacter acanthamoeba (*12*), endosymbiont of *Acanthamoeba* UWC8 (Wang *et al* 2014), *Occidentia massiliensis* Os18 (*79*), Rickettsiales bacterium Ac37b and alphaproteobacterial outgroups *Caulobacter crescentus* CB15, *Candidatus* Puniceispirillum marinum IMCC1322 (*80*), *Azospirillum brasiliense* Sp245 (now *Azospirillum baldaniorum*) (*81*), MarineAlpha3 Bin5, MarineAlpha3 Bin2, MarineAlpha12 Bin1, MarineAlpha11 Bin1, MarineAlpha9 Bin6 and MarineAlpha10 Bin2 (*14*). Orthologs were identified through PSI-BLAST searches using the 129 gene alignments as a query and non-orthologs were detected and removed via single gene tree inspections as described above. A discordance filter (*82*) was applied to remove the most discordant genes: (i) single gene alignments were prepared with MAFFT E-INS-i (*74*) and trimAl -gappyout (*75*), (ii) single gene trees were inferred with IQTREE v.1.6.9 (*50*) (*83*) (with -bnni), (iii) bipartition count profiles were constructed from the bootstraps (tre_make_splits.pl) and compared between all possible gene pairs to calculate discordance scores (tre_discordance_two.pl). The top 13 most discordant genes were removed from the dataset (Figure S3). After preliminary phylogenomics analyses with the remaining 116 genes, we decided to omit extremely long branching taxa *Candidatus* Xenolissoclinum pacificiensis and *Candidatus* Fokinia solitaria and phylogenetic unstable taxa endosymbiont of *Stachyamoeba lipophora* and UBA6177 from downstream phylogenomics analyses (Data S1). The resultant 116 orthologous groups were first aligned by applying PREQUAL v1.01 (*84*) (masking of putative non-homologous sites), MAFFT E-INS-i (multiple sequence alignment) and Divvier -partial (*85*) (alignment ‘divvying’) and then concatenated into an “untreated” supermatrix alignment. From this, an iterative χ^2^-trimmed alignment (47 rounds of removing the top 1% most heterogeneous sites as determined by χ^2^-score (See *48*) was prepared. The iterative χ^2^-trimmer is more efficient at reducing compositional heterogeneity compared to the standard χ^2^-trim method (*14, 86*). These were used for phylogenetic reconstruction under the CAT+GTR+Γ4 (untreated) and CAT+LG+Γ4 (iterative χ^2^-trimmed) model with PhyloBayes MPI v1.8 (*87*). Four independent MCMC chains were run until convergence was reached (maxdiff < 0.3) or a sufficient effective sample size was reached (effsize > 300), while using a burnin of at least 5000 generations. Posterior predictive checks were performed to check to what degree the inferred phylogenetic models captured the across-taxa compositional heterogeneity and site-specific pattern diversity present in the alignments. Parameter configurations were sampled every 50 generations after the burnin. Maximum likelihood phylogenetic reconstructions were done under the PMSF approximation (with 100 non-parametric bootstraps; guidetree under LG+G+F) of the LG+C60+F+Γ4 model for both supermatrix alignments with IQTREE v1.6.5.

#### Gene family trees

We used the annotation of AlphaNOGs from EggNOG 4.5.1 to assign proteins from the MAGs and reference genomes into clusters. All proteins without AlphaNOG annotation were subjected to all-versus-all BLASTP analysis and subsequent de-novo clustering with Silix (overlap 90, identity 60) (*88*), resulting in a total of 34361 clusters. We computed alignments for all protein clusters with at least 4 members (4240), using PREQUAL (*84*) to prefilter unaligned sequences, MAFFT E-INS-i to perform alignments and Divvier (-divvygap) (*85*) to filter alignments. Single gene trees were inferred for the 4240 alignments with IQTREE v1.6.9 (*50*) with 1000 ultra-fast bootstraps (-bb 1000 -wbtl) (*89*). A model test was performed (-m TESTNEW -mset LG -madd LG+C10,..,LG+C60) for each tree inference (*83*). For clusters with only 2 or 3 members we created ‘dummy’ bootstraps that represent the only possible topology for an unrooted tree.

#### Gene tree-species tree reconciliation

The 4240 single gene trees were reconciled with the species tree derived from the iterative χ^2^-trimmed alignment with CAT+LG+Γ4 using the ALEml_undated algorithm of the ALE suite (*20*). ALE infers gene duplications, losses, transfers, originations and ancestral gene contents along a species tree. Per node, a gene family was considered present if ‘copies’ ≥ 0.3 in the ALE output. In addition, gene family evolution events (transfers, originations and losses) were counted similarly with a threshold of 0.3. This threshold ensures that events that were reconstructed with a low frequency are still detected, since the signal in single gene trees can sometimes be very low and events could be overlooked at more stringent thresholds (*86*, See also *90*). Singleton clusters were counted as originations for the corresponding species.

#### Environmental diversity

The 16S rRNA gene sequences of the MAGs (Gamibacteraceae and Mitibacteraceae, no 16S sequences were found for Athabascaceae) were used to search the NCBI nt database with BLASTN (E-value <0.05, length > 700 bp). An alignment was prepared for Rickettsiales and an alphaproteobacterial outgroup using MAFFT E-INS-i, and trimmed with trimal (-automated1). A phylogenetic tree was inferred with IQTREE with 100 nonparametric bootstraps under the GTR+F+R8 model (selected by IQTREE’s model test).

#### T4SS subunit gene trees

Trees for the T4SS subunits virB1-6,8-11,D4 were prepared using the corresponding eggNOG clusters for all Bacteria. First, we removed all Rickettsiales sequences from these clusters in order to avoid self-alignment. We then used the sequences from our taxon selection as queries to search the COGs using DIAMOND (--top 50 --ultra-sensitive) and clustered the obtained sequences using cd-hit at 80% identity. We then aligned those reference sequences together with the sequences from our taxon sampling using MAFFT-E-INS-i and applied a light gap trimming (trimal -gt 0.01). Finally trees were inferred in IQ-TREE with automatic model selection and support values were estimated by 1000 ultrafast bootstraps.

#### Ancestral gene content reconstruction

We reconstructed ancestral gene family repertoires from ALE by selecting all gene families predicted to be present at a given node with a frequency ≥ 0.3. Consensus annotations for gene families were inferred based on the abovementioned gene annotations. Enzyme complexes were inferred as present if at least half of the necessary genes were present. We assessed the metabolic capabilities of ancestral genomes using the KEGG Module tool (*60*) or MetaCyc pathways (*91*).

### Supplementary Text

#### Taxonomic Descriptions

##### Description of “Candidatus Mitibacter marchionensis” (sp. nov, gen. nov.)

*Miti* translates to “Sea” in Tahitian, close to the Marquesas Islands (latinised as marchionis), with suffix -ensis. Exclusively marine sequences, larger genomes than other Rickettsiales, flagellar and chemotaxis genes present, larger metabolic capacity (e.g. genetic repertoire for biosynthesis of all amino acids, nucleotides, etc.). 16S and 23S available.

##### Description of Candidatus Mitibacteraceae (fam. nov.)

Description is the same as for the genus Mitibacter, except that several MAGs were isolated from freshwater (UBA6178, UBA6189, UBA6149) (*42, 43*). Suff. -aceae, ending to denote a family. Type genus: Mitibacter

##### Description of Candidatus Athabascaceae (fam. nov.)

Prefix Athabasc-from the sampling location, the Athabasca oil sands syncrude tailing pond surface water near Fort McMurray in northeastern Alberta, Canada, from which the most complete MAG UBA6187 was extracted (*42, 92*). Also contains marine sequences (TARA_ANE_MAG_00011) (*44*). Similarly to Mitibacteraceae, larger genomes than other Rickettsiales, flagellar and chemotaxis genes present, larger metabolic capacity (e.g. genetic repertoire for biosynthesis of all amino acids, nucleotides, etc.). No rRNA available. Suff. -aceae, ending to denote a family.

##### Description of Candidatus Gamibacteraceae (fam. nov.)

The word “Water” translates to //gam-i in Khoekhoe, a language of the Khoe languages, which are spoken by San and Khoekhoen people, native to South Africa. The MAGs C-132 and C-137 were both extracted from a marine sample that was taken close to the Cape region. Other MAGs were isolated from freshwater (UBA2645) and an aquifer (Alphaproteobacteria bacterium RIFCSPLOWO2 01 FULL 40 26) (*42, 45*). Reduced metabolic capacity (pathways for 18 amino acids present, nucleotide metabolism present), no flagellar or chemotaxis genes and no Tad system, no ATP/ADP translocase. 16S and 23S available. Suff. -aceae, ending to denote a family.

#### Phylogeny of the Gamibacteraceae and Deianireaeceae

The Alphaproteobacteria, of which the Rickettsiales are part of, are particularly sensitive to phylogenetic artefacts (*14, 47, 48, 72*). The generally AT-rich Rickettsiales often branch artifactually with other AT-rich alphaproteobacterial lineages such as Pelagibacterales, alpha proteobacterium HIMB59, MarineAlpha6, -7, -8 and -9 and the Holosporaceae. Although the phylogenetic relationships within the Rickettsiales are generally quite stable in the literature, there is some disagreement with respect to the placement of the Midichloriaceae. In some studies, they branch as sister to Rickettsiaceae (*93*), while in others they branch as sister to the Anaplasmataceae (*94, 95*).

To test for the possible influence of compositional bias artefacts on the expanded Rickettsiales species tree, we inferred maximum likelihood and Bayesian inference phylogenies from an “untreated” supermatrix alignment and one that had the most heterogeneous sites removed via an iterative χ^2^-trimming approach. The resultant trees were highly congruent, with the exception of the placement of the Gamibacteraceae and Deianireaeceae. In the “untreated” trees, they branch as a clade sister to the Anaplasmataceae with near-maximum-to-maximum branch support (99 UFB, 100 NPB, 1.00 PP) and in the “iterative χ^2^-trimmed” trees, they branch as either unresolved with respect to Rickettsiaceae, Midichloriaceae and Anaplasmataceae (50 UFB, 53 NPB) or as sister to a clade comprising Midichloriaceae and Anaplasmataceae with near maximum branch support (0.99 PP). This suggests that the placement of the Gamibacteraceae and Deianireaeceae clade sister to Anaplasmataceae (also observed in (*11*)) may be the result of a phylogenetic artefact.

#### Putative T4SS effectors in free-living Rickettsiales

While no Rickettsiales effector proteins seem to be conserved across families, many of the experimentally verified effectors contain eukaryotic-like repeat domains such as Ankyrin and LRR (*7, 12, 96*). We found no enrichment of such domains in Mitibacteraceae (and, to a lesser degree, in Athabascaceae) compared to other free-living alphaproteobacteria. Instead, we could identify several domains that have been identified in effector proteins of the bacteria-killing T4SS of *Xanthomonas citri*, such as pepdidoglycan binding domains (PGBD) and Peptidase M23 (*28, 29*).

#### Key transitions in classical Rickettsiales family ancestors

Our reconstructions infer gene loss, i.e., reductive genome evolution, as the major evolutionary event in ancestral Rickettsiales diversification, as expected for obligate host-associated organisms (*4*). While considerably smaller than its free-living ancestor, our reconstructions suggest a larger genome in LhRCA (1,165 genes) than all family rank ancestors of classical Rickettsiales (734-1,147 genes; Fig. S12; Data S5). Reduction of metabolic capabilities is typical for specialized endosymbionts (*97*) and strikingly occurred independently at least twice during host-associated Rickettsiales evolution. Severe reduction in amino acid biosynthesis affected the common ancestor of Midichloriaceae and Anaplasmataceae (LMiACA), as well as Rickettsiaceae (LRiCA), losing the potential to synthesize nine and eleven amino acids (Fig. S12), respectively. These lineages contain the well-known animal pathogens and highly specialized insect symbionts and this independent pattern of metabolic gene loss underpins that the evolution of animal association occurred twice in Rickettsiales. Both ancestors, however, gained genes associated with their specialized animal-associated lifestyle. LRiCA gained key genes involved in host entry and manipulation, i.e. rickettsial outer membrane protein and ankyrin repeat containing effectors (Fig. S12; Data S6), respectively. LMiACA, on the other hand, gained an additional DnaJ-like chaperone, likely aiding to compensate for the accumulation of slightly deleterious mutations associated with relaxed selection and large genetic drift in animal endosymbionts (*98*). The second notable gain in LMiACA represents a DsbB protein that has been proposed to be involved in oxidative folding of secreted *Wolbachia* effectors (*99*).

Over Rickettsiales evolution, independent loss also affected the ancestral potential to store carbon as polyhydroxybutyrate (PHB) (Fig. S9; Fig. S12; Data S6). This common trait of many marine heterotrophic bacteria (*100*) and several representatives of Rickettsiaceae (*101*) was lost in the last common ancestor of the Gamibacteraceae and *Deianiraea vastatrix* (LGDCA) as, well as the Anaplasmataceae (LACA). Intriguingly, losing the potential to synthesize the storage compound PHB in both ancestors co-occurred with the loss of flagellar biosynthesis (Fig. S12) and could indicate a diminished capability for host-free motility and survival in LACA and LGDCA.

The hallmark feature of energy parasitism, the ATP/ADP translocase, was lost twice in Rickettsiales evolution in the last common ancestor of the Gamibacteraceae (LGCA) and LACA. It is known that Anaplasmataceae produce their own ATP via oxidative phosphorylation (*1*) and this potential is likewise encoded in Gamibacteraceae MAGs (Fig. S9). The genes for adhesin production and export (*fha*C, *fha*B) were gained in the LGDCA and were proposed to be involved in host cell adhesion of the episymbiont *Ca*. Deianiraea vastatrix (*11*) and was shown to be essential for host cell adhesion in *Bordetella* (*102*). Their conservation in *Ca*. Deianiraea vastatrix and Gamibacteraceae MAGs underpins a likely ancestral ectosymbiotic lifestyle of this group. *Ca*. Deianiraea vastatrix is auxotrophic for multiple amino acids and likely employing energy parasitism, while Gamibacteraceae show larger metabolic capabilities and are likely less host dependent for energy and nutrients (Fig. 3; Fig S9).

---

**Fig. S1.**
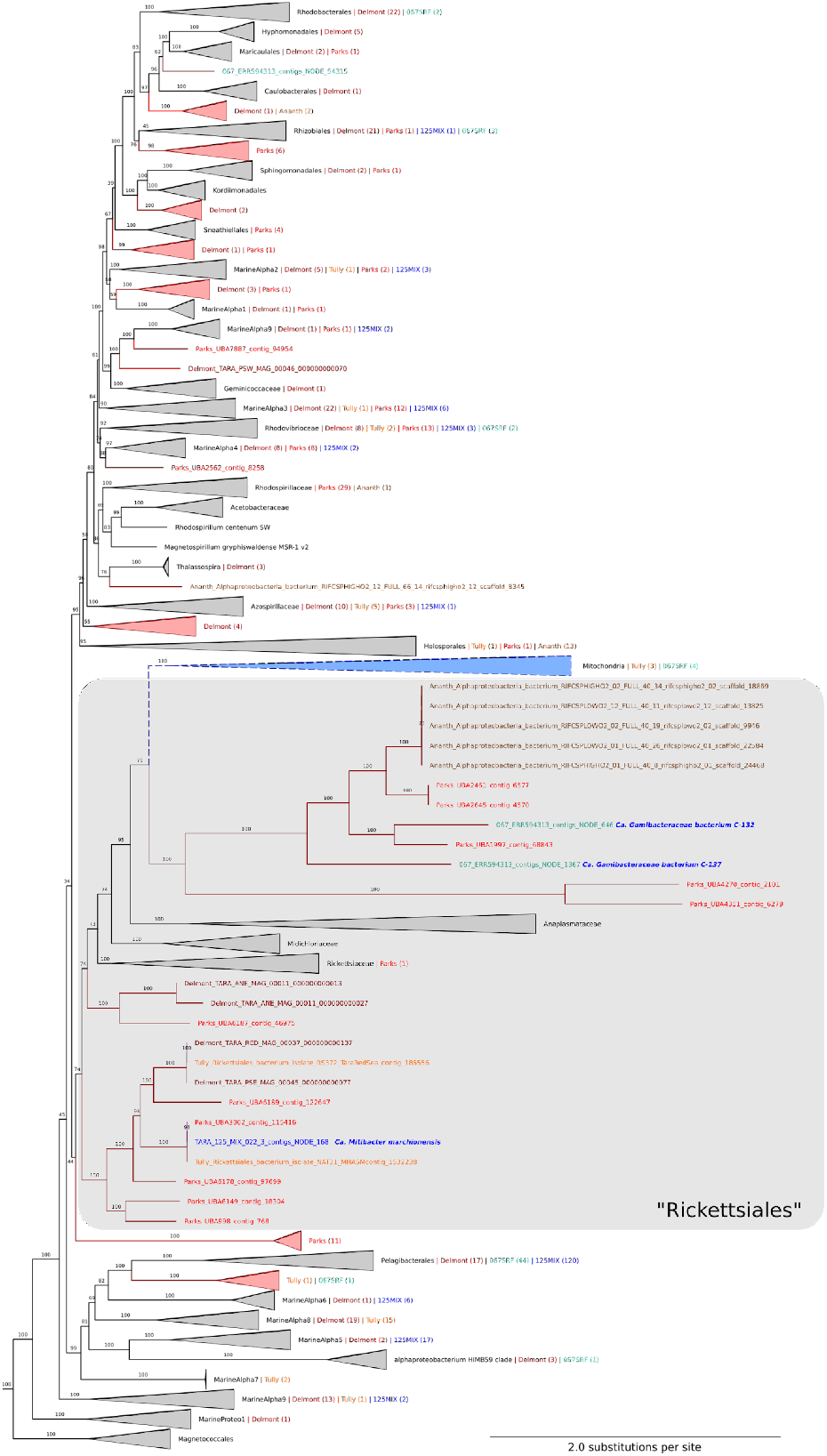
Alphaproteobacterial and mitochondrial phylogenetic diversity in analysed metagenomic datasets. Phylogenetic analyses of concatenated ribosomal protein sequences (see Methods) identified in the metagenome assemblies of the two Tara Ocean’s environmental samples ‘TARA_125_MIX_022-3’ (dark blue) and ‘TARA_067_SRF_0.22-0.45’ (teal) and the metagenome assembled genomes of Tully *et al* (*43*) (orange), Parks *et al* (*42*) (red), Delmont *et al* (*44*) (brown-red) and Anantharaman *et al* (*45*) (light-brown). Reference taxa are given in black. Clades are collapsed in grey triangles when they include reference taxa and in red triangles when they include no reference taxa. The number of environmental alphaproteobacteria and mitochondria is denoted in the collapsed triangle annotation. Length of mitochondrial stem branch is shortened to improve tree readability. Tree was reconstructed with the RP15 pipeline (see Methods). The complete tree is available as a Newick file (Data S1).

**Fig. S2.**
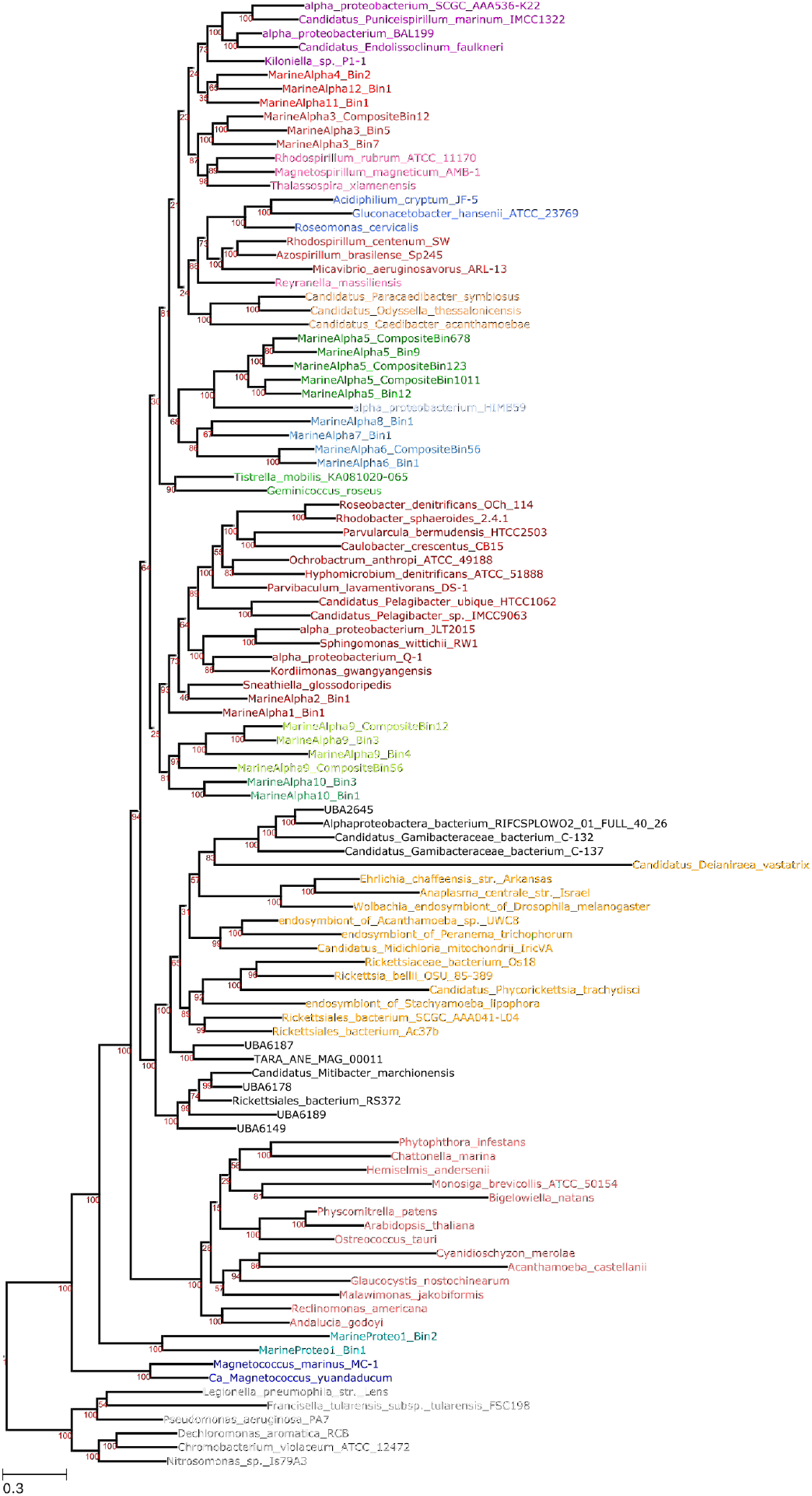
Maximum likelihood phylogeny of Alphaproteobacteria and mitochondria. Based on the χ^2^-trimmed (20% most heterogeneous sites removed) concatenated alignment of 24 highly conserved mitochondrially encoded proteins. ML tree was inferred under the PMSF approximation of LG+C60+F+Γ4 with 100 non-parametric bootstraps as implemented by IQ-TREE. Distinct alphaproteobacterial clades are given their own unique color, including the Rickettsiales (orange) and mitochondria (salmon). Novel Rickettsiales MAGs identified and/or reconstructed in this study are given in black.

**Fig. S3.**
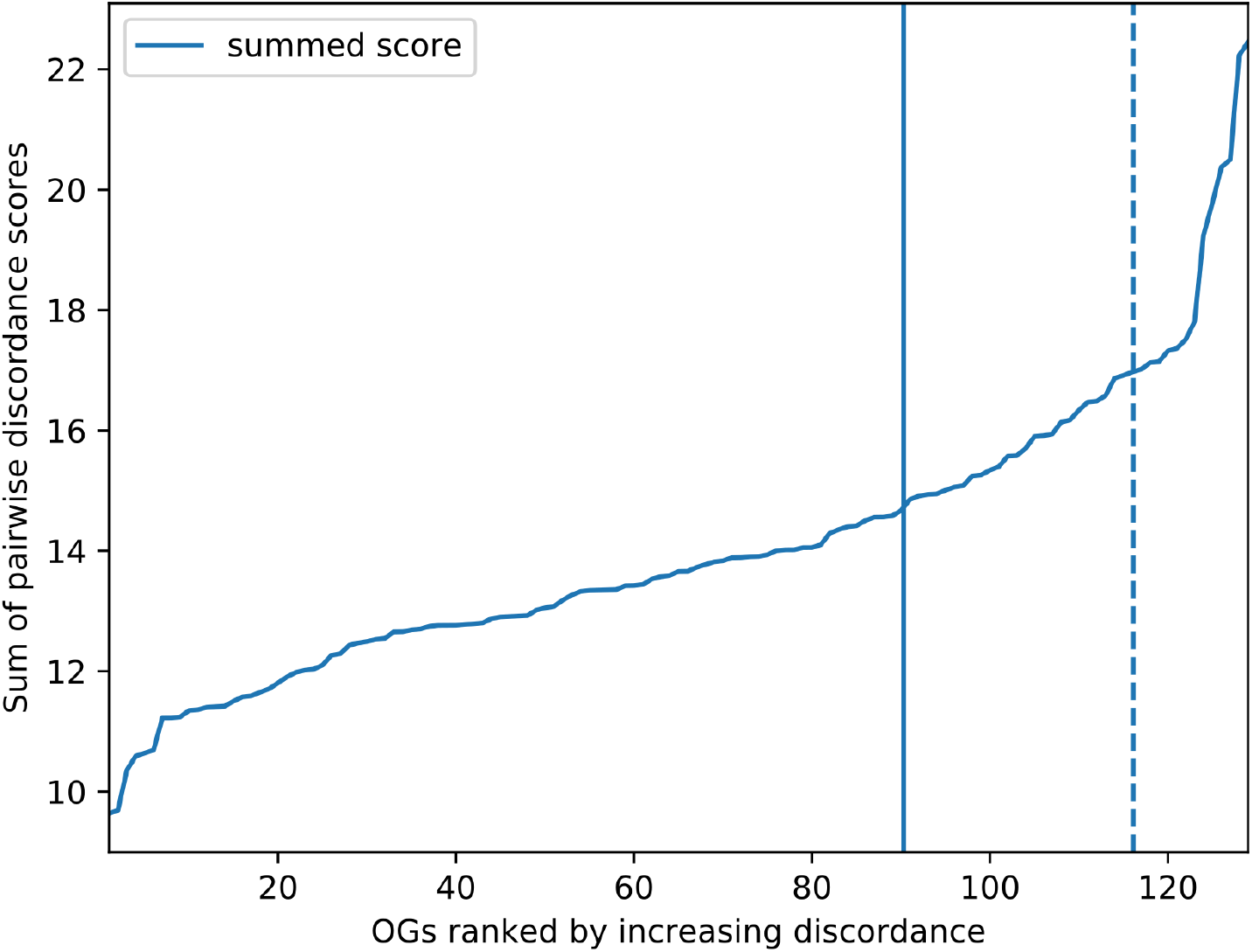
Discordance filter of the 129 panortholog dataset from Martijn et al. (*35*). Bipartition count profiles were constructed from the bootstraps of each single gene tree and compared between all possible gene pairs to calculate discordance scores. Genes were then ordered according to increasing discordance. The 13 most discordant genes, scoring above the dotted line, were rejected.

**Fig. S4.**
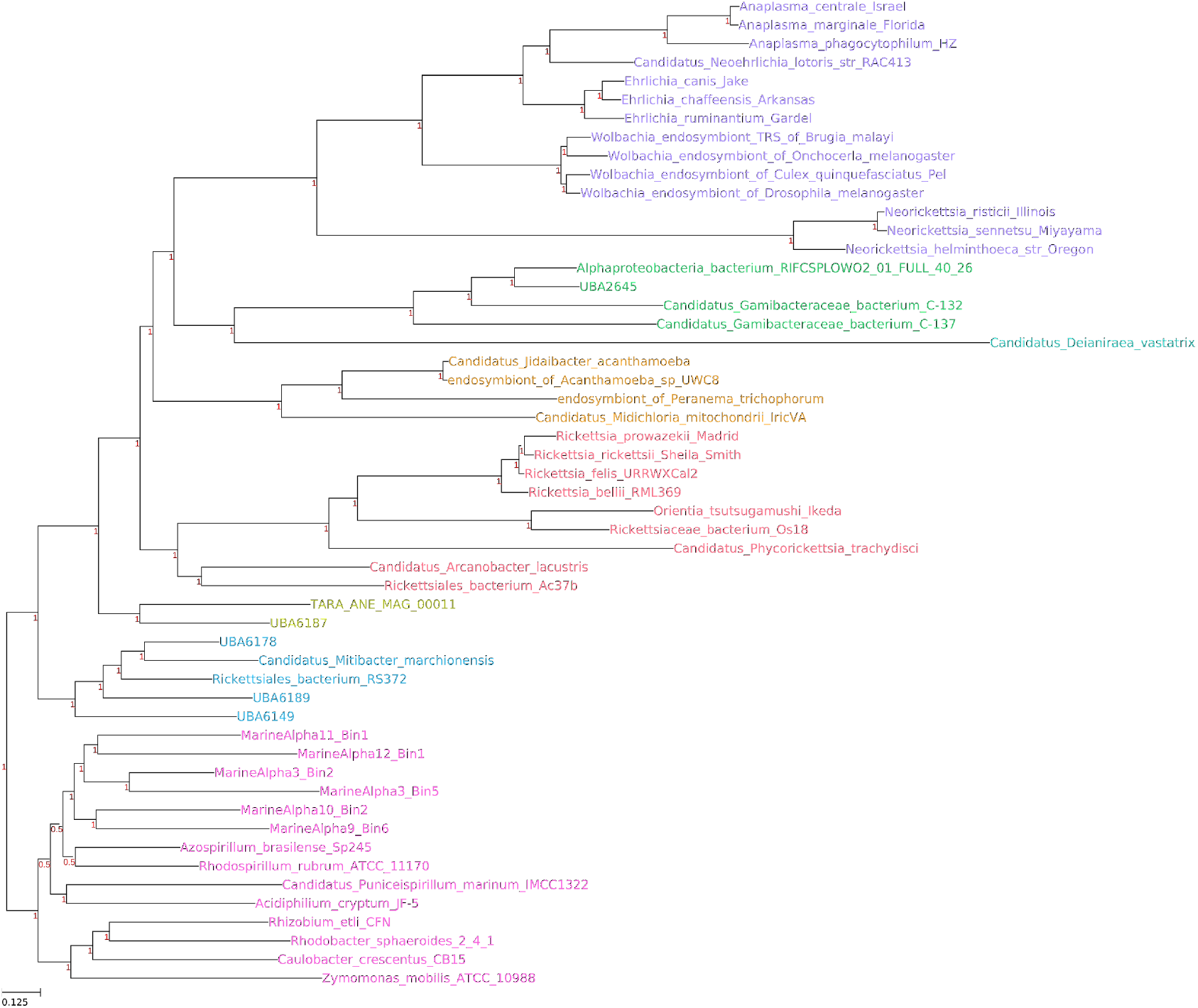
Bayesian inference of Rickettsiales phylogeny based on the untreated concatenated alignment of 116 panorthologs. Consensus tree over all 4 MCMC chains that were inferred under the CAT+GTR+Γ4 model is given. Anaplasmataceae (purple), Gamibacteraceae (green), Deianiraea (teal), Midichloriaceae (orange), Rickettsiaceae (salmon), Athabascaceae (moss), Mitibacteraceae (blue) and other alphaproteobacteria (pink). The four trees converged with respect to the in-group, but failed to converge with respect to the outgroups. Consensus trees of the individual MCMC chains are available as Newick files (Data 1).

**Fig. S5.**
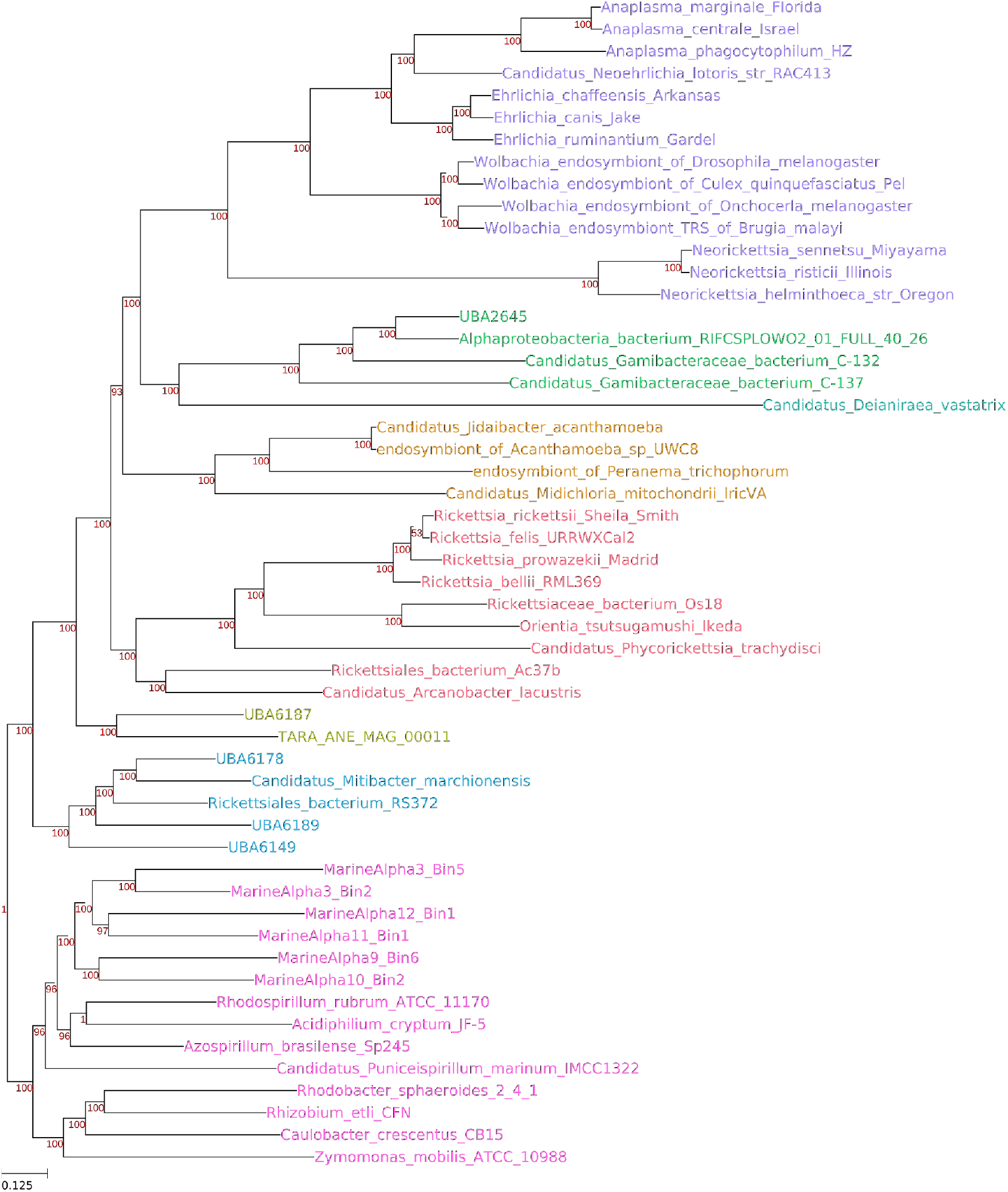
Maximum likelihood phylogeny of Rickettsiales based on the untreated concatenated alignment of 116 panorthologs. ML tree inferred under the PMSF approximation of LG+C60+F+Γ4 with 100 non-parametric bootstraps as implemented by IQTREE. Anaplasmataceae (purple), Gamibacteraceae (green), Deianiraea (teal), Midichloriaceae (orange), Rickettsiaceae (salmon), Athabascaceae (moss), Mitibacteraceae (blue) and other alphaproteobacteria (pink).

**Fig. S6.**
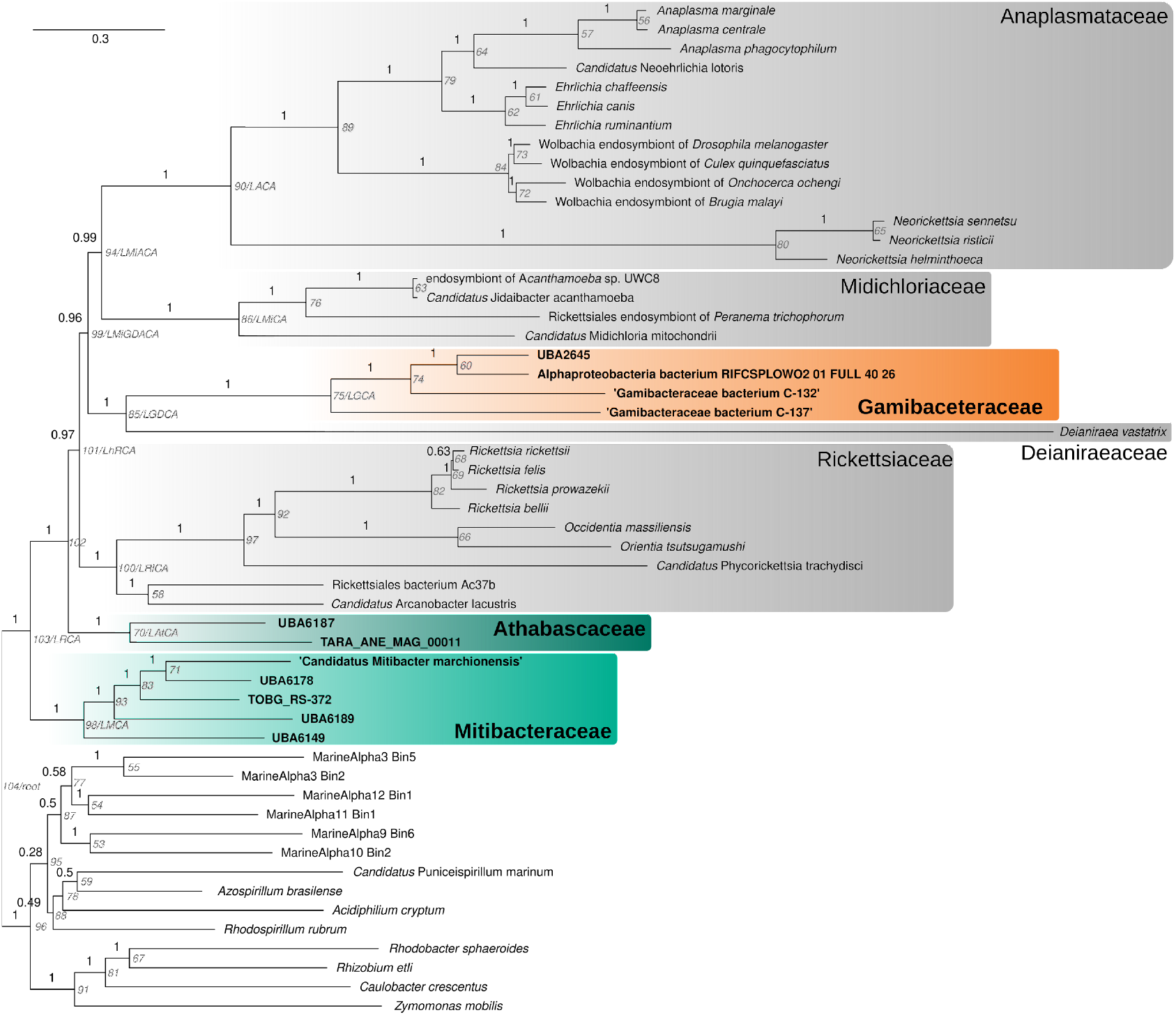
Bayesian inference Rickettsiales species tree with alphaproteobacterial outgroup based on a dataset of 116 marker genes. The alignment was prepared using iterative χ^2^-trimming: 47 rounds of removing the top 1% most heterogeneous sites as determined by χ^2^-score), highlighting the phylogenetic position of Gamibacteraceae (orange shading), Athabascaceae (dark green shading) and Mitibacteraceae (light green shading). The tree was inferred with PhyloBayes (CAT+LG+Γ4, 20000 cycles with a burn-in of 5000 cycles). Support values (in black) correspond to posterior probabilities. Internal node names and ancestor designations used for the ancestral reconstruction are written in grey. Last common ancestor of Anaplasmataceae, Midichloriaceae, Deianiraea and Gamibacteraceae: LMiGDACA; Last common ancestor of Anaplasmataceae and Midichloriaceae: LMiACA; Last common ancestor of Deianiraea and Gamibacteraceae: LGDCA; Last common ancestor of Anaplasmataceae: LACA, Midichloriaceae : LMiCA, Gamibacteraceae: LGCA; Rickettsiaceae: LRiCA, Mitibacteraceae: LMCA, Athabascaceae: LAtCA.

**Fig. S7.**
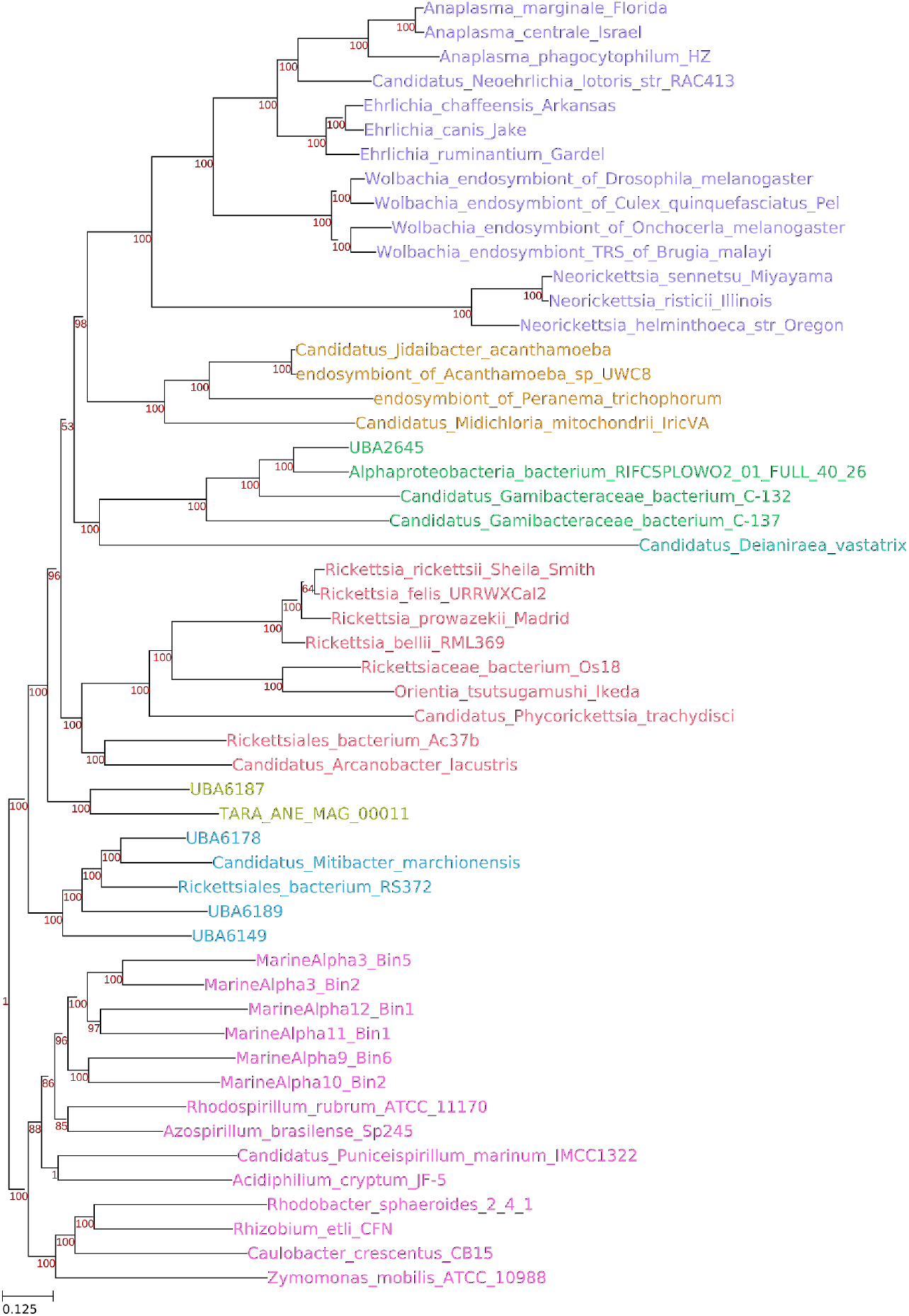
Maximum likelihood Rickettsiales species tree with alphaproteobacterial outgroup based on a dataset of 116 marker genes. The alignment was prepared using iterative χ^2^-trimming: 47 rounds of removing the top 1% most heterogeneous sites as determined by χ^2^-score. ML tree inferred under the PMSF approximation of LG+C60+F+Γ4 with 100 non-parametric bootstraps as implemented by IQTREE. Anaplasmataceae (purple), Gamibacteraceae (green), Deianiraea (teal), Midichloriaceae (orange), Rickettsiaceae (salmon), Athabascaceae (moss), Mitibacteraceae (blue) and other alphaproteobacteria (pink).

**Fig. S8.**
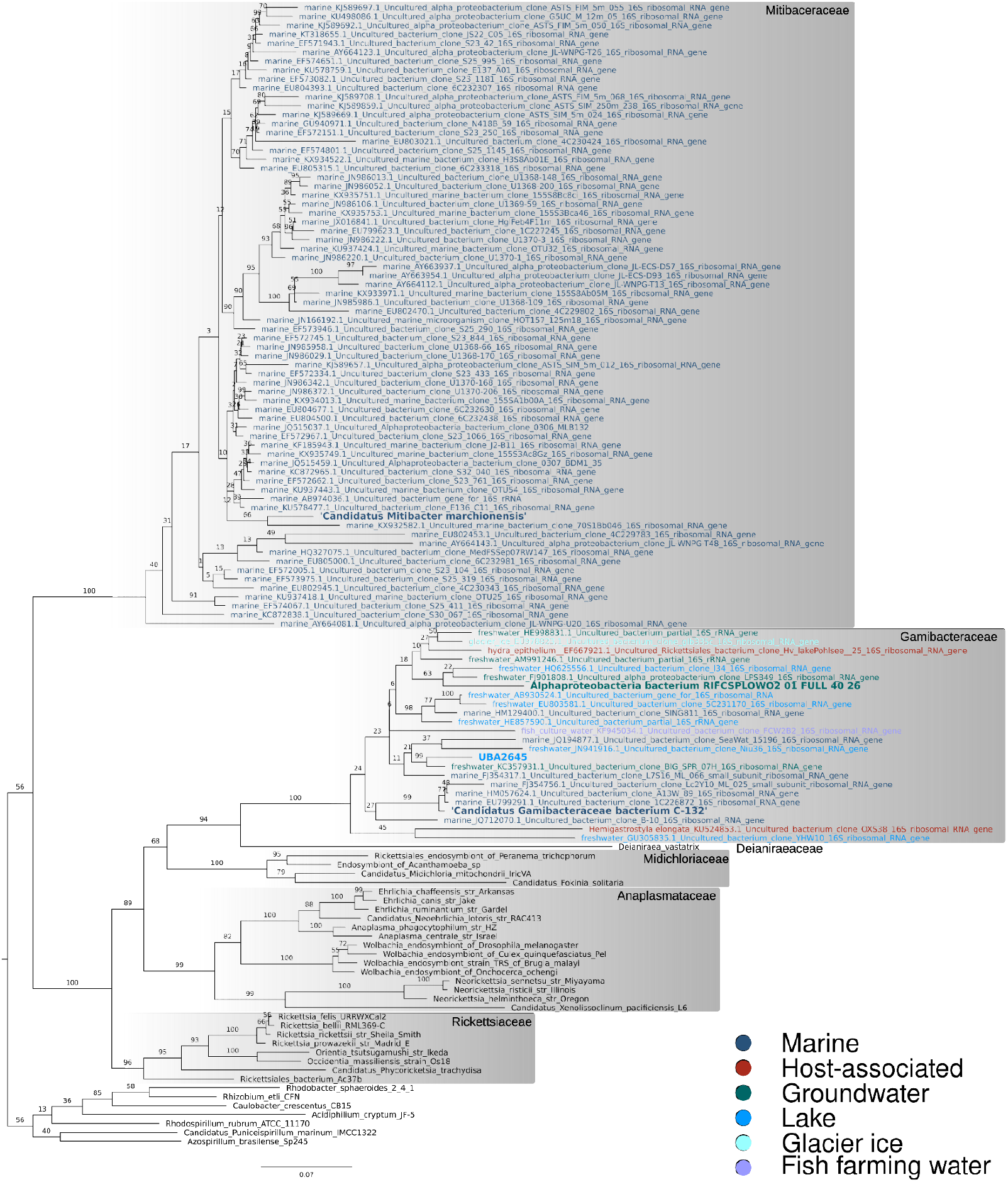
Phylogenetic tree of 16S rRNA gene sequences related to Mitibacteraceae and Gamibacteraceae. Sequences of environmental origin are color-coded (see legend) and reference sequences from other Rickettsiales families and an alphaproteobacterial outgroup are indicated. Support values correspond to 100 non-parametric bootstraps. The tree was reconstructed in IQ-Tree under the GTR+F+R8 model (see Methods).

**Fig. S9.**
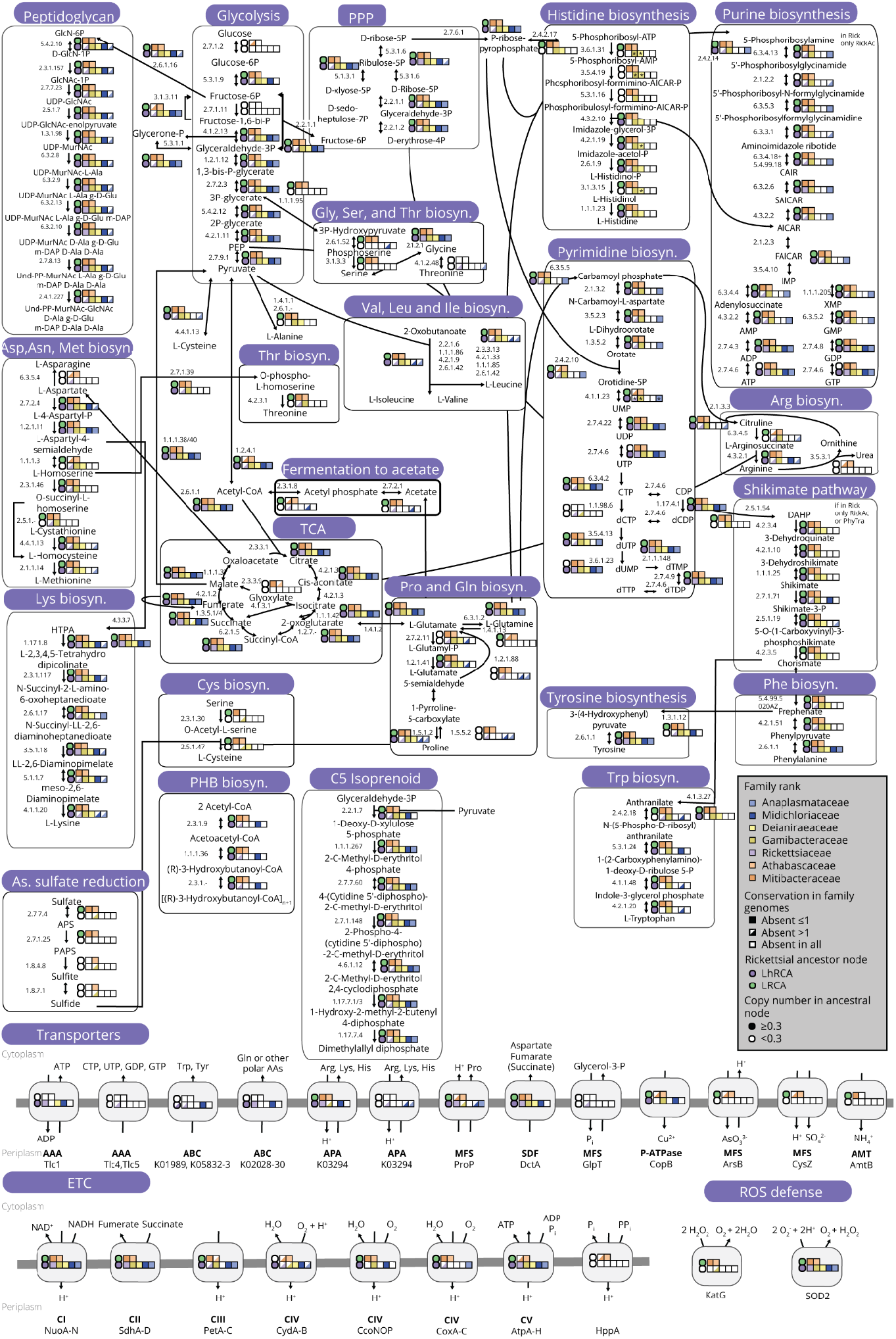
Overview of conservation of central metabolism, transporters, and electron transport chain (ETC) in Rickettsiales families and key ancestors. All enzymatic steps are depicted with enzyme commision numbers. The corresponding enzyme name and distribution on Rickettsiales number can be found in Data S6. Arrows between molecules and arrow heads indicate catalyzed enzymatic reactions and directionality, respectively.

**Fig. S10.**
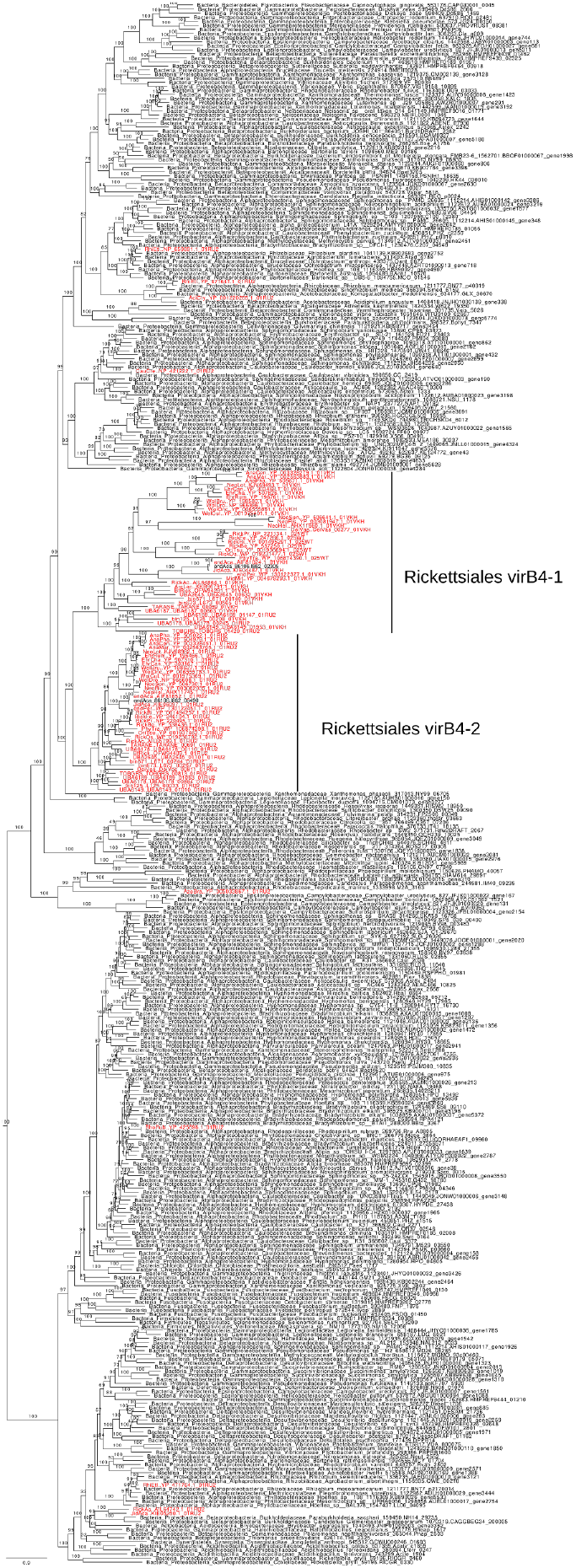
Phylogenetic tree of virB4 homologs from the sampled Rickettsiales genomes and reference sequences from the EggNOG v5.0 database (COG3451). Tree was reconstructed in IQ-TREE under the model LG+F+R10 with 1000 ultrafast bootstraps. Taxa included in the ancestral reconstruction are highlighted in red, reference sequences include a taxonomic label. Rickettsiales virB4-1 and virB4-2 are monophyletic, suggesting a common duplication in LRCA.

**Fig. S11.**
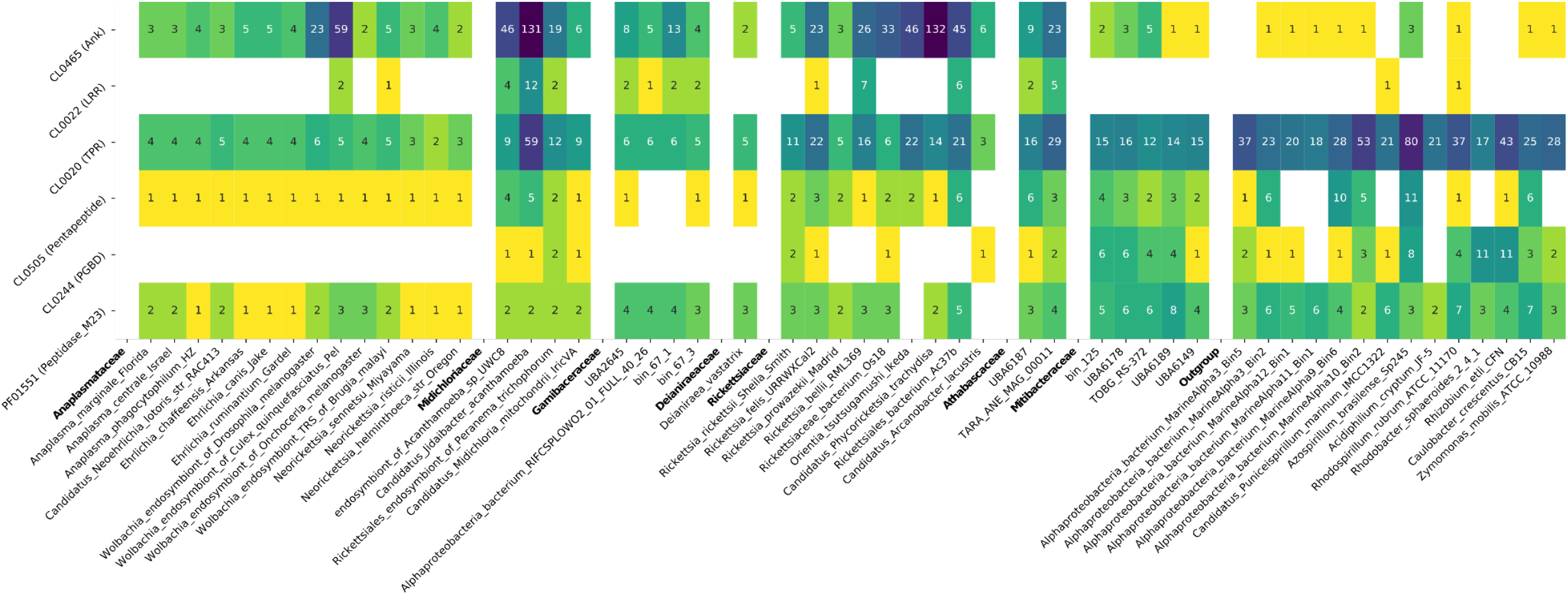
Frequency of proteins containing selected PFAM domain families. Domains are typical of Rickettsiales effectors (Ank, LRR, TPR) and domains often found in Xanthomonas bacterial-killing effectors (Peptidoglycan binding domain, Peptidase M23) in the selected taxa used for phylogenomic and ancestral reconstruction. Domains were identified using HMMER hmmsearch -E 1e-05.

**Fig. S12.**
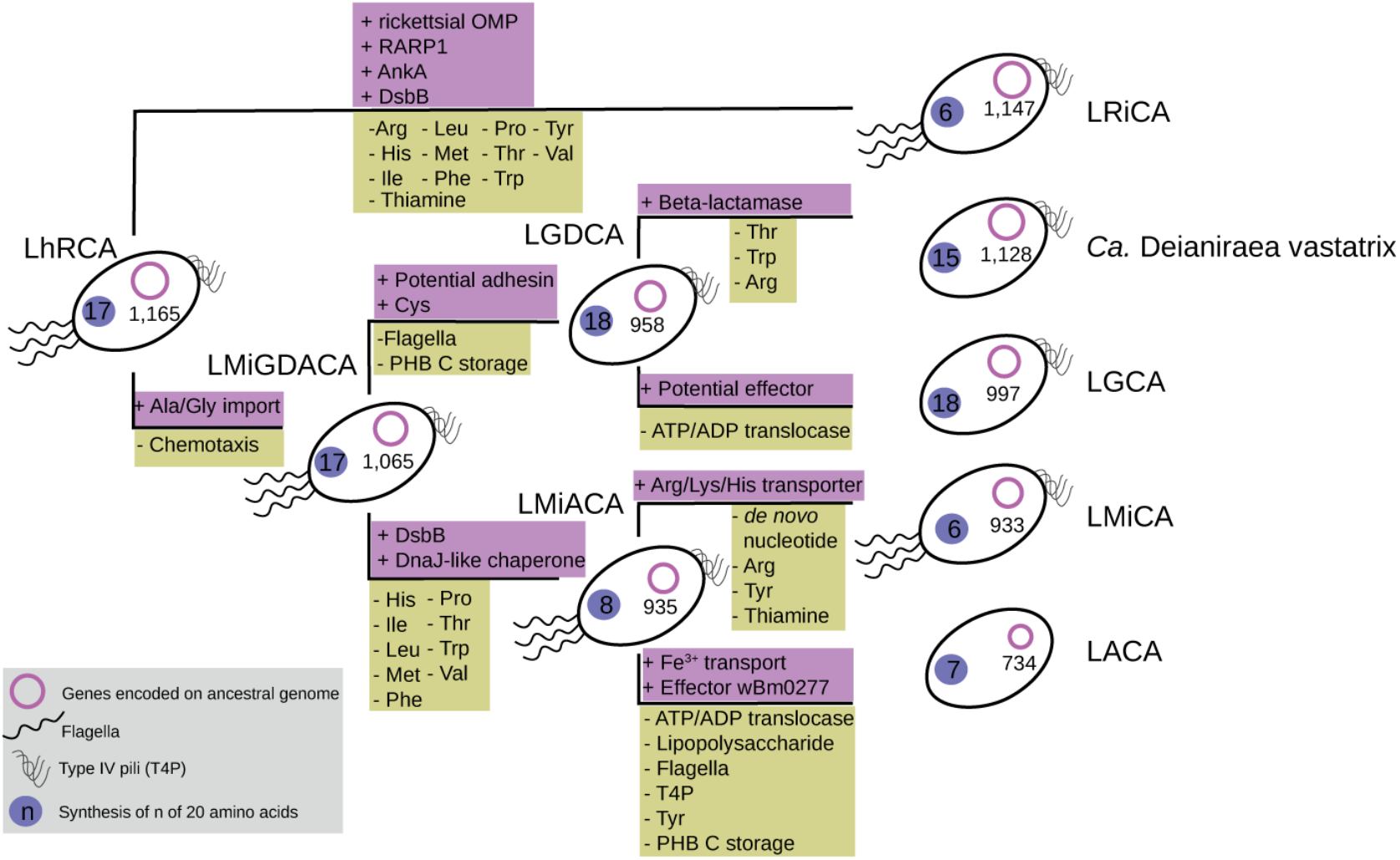
Key gains, losses and characteristics of host-associated Rickettsiales ancestors. Schematic phylogenetic tree depicting the relationships of the last common ancestor of host-associated Rickettsiales (LhRCA) and Rickettsiales families ancestors. Last common ancestor of Anaplasmataceae, Midichloriaceae, Deianiraea and Gamibacteraceae: LMiGDACA; Last common ancestor of Anaplasmataceae and Midichloriaceae: LMiACA; Last common ancestor of Anaplasmataceae: LACA, Midichloriaceae: LMiCA, Gamibacteraceae: LGCA; Rickettsiaceae: LRiCA. Gains (violet) and losses (green) of genes encoding for key characteristics are written above and below the corresponding branch, respectively. The ancestors are depicted as cells with inferred features such as the presence of a flagellum, Type 4 pili, the capacity to synthesize amino acids and the number of inferred ancestral genes. Amino acid biosynthesis pathways are represented by the three-letter code of the produced amino acid. Proteins related to ankyrin-repeat containing protein (AnkA); disulfide bond formation (DsbB); rickettsial outer membrane protein (RARP); polyhydroxybutanoate (PHB) synthesis.

**Fig. S13.**
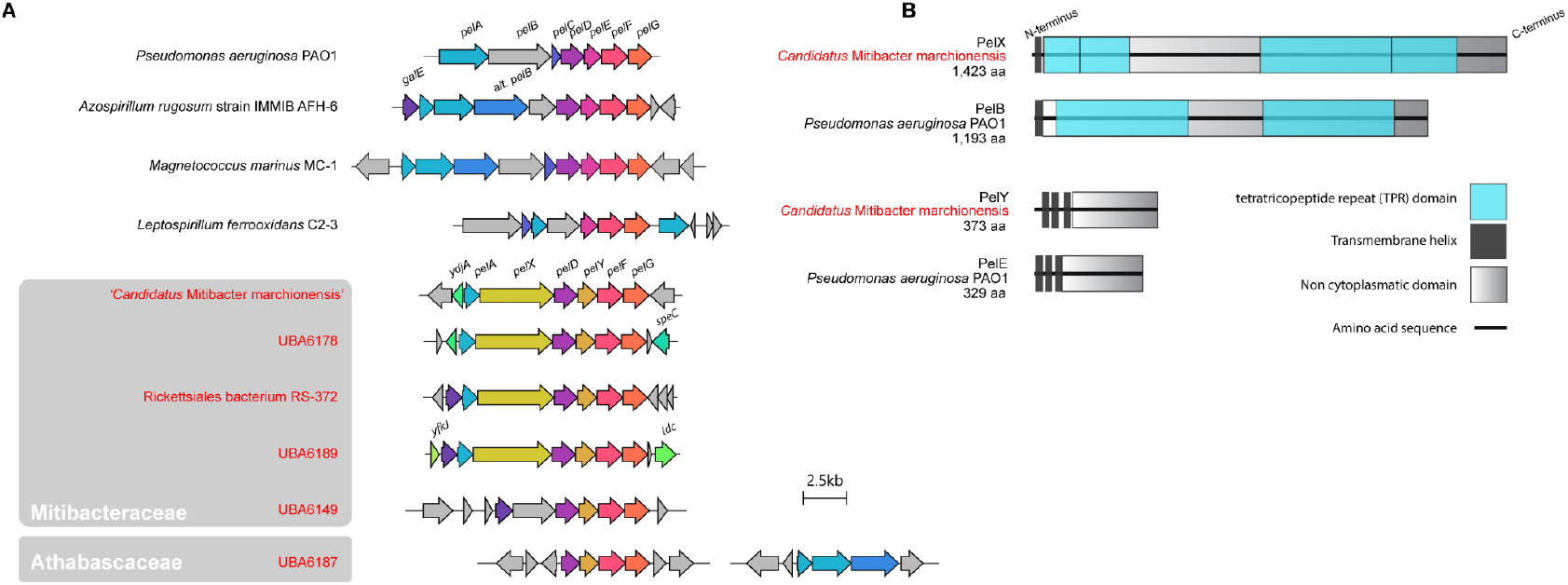
Gene synteny of and domain architecture of derived pel exopolysaccharide gene cluster in free-living Rickettsiales. (A) Gene synteny plot of *pel*ABCDEFG gene cluster in the model organism *Pseudomonas aeruginosa* PAO, and related clusters in selected free-living bacteria, and free-living Rickettsiales. Arrows indicate the genomic orientation and size of genes. Scale bar indicates a length of 2.5 kb. (B) Domain architecture of likely functional equivalents of the *P. aeruginosa* PelB and PelE proteins in the Rickettsiales bacterium *Candidatus* Mitibacter marchionensis.

**Table S1.** Overview of Bayesian phylogenetic analyses. Summary statistics of alignments, MCMC sampling and posterior predictive tests of Bayesian phylogenetic analyses.

| **Dataset: 116 near-panorthologs Rickettsiales** | **Untreated** | **iterative $\chi^2$-trim**<br>**(47 rounds of**<br>**removing the top**<br>**1% sites)** |
| --- | --- | --- |
| Sites | 38842 | 24238 |
| Informative sites | 29598 | 15347 |
| Missing data (%) | 21.3 | 24.1 |
| $\chi^2$-score | 78360 | 423 |
| Model of evolution | CAT+GTR+Γ4 | CAT+LG+Γ4 |
| *MCMC sampling statistics* |  |  |
| Generations | 32000 | 20100 |
| Burnin | 15000 | 5000 |
| Effective sample size* | >533 | >240 |
| Maxdiff | 1 | 1 |
| *Posterior predictive tests\*\** |  |  |
| Maximum squared heterogeneity (p-value) | 0 | 0.87 - 0.90 |
| Maximum squared heterogeneity (Z-score) | 240 - 278 | -1.1 - -1.1 |
| Mean squared heterogeneity (p-value) | 0 | 0.90 - 0.94 |
| Mean squared heterogeneity (Z-score) | 966 - 1075 | -1.3 - -1.4 |
| Mean diversity per site (p-value) | 0.01 - 0.04 | 0.04 - 0.06 |
| Mean diversity per site (Z-score) | 2.0 - 2.1 | 1.5 - 1.7 |
| * Considering only log-likelihood, total tree length, alpha parameter and number of categories |  |  |
| ** Range across 4 chains |  |  |

**Data S1. (separate file)**

**Supplementary phylogenetic trees**. Tar archive with all phylogenetic trees (rp15, alphamito24 and the 116 gene dataset) showing the position of the novel MAGs in newick format.

**Data S2. (separate file)**

**Sequencing runs of samples used for differential coverage binning**. Samples that were assembled and of which the contigs were binned are highlighted in orange. All samples were used for differential coverage binning of TARA_125_SRF_0.22-3. Samples highlighted in green and ERR594313 were used for differential coverage binning of TARA_067_SRF_0.22-0.45. This table is a subset of the companion Table W1 published by Sunagawa et al, 2015 (*37*).

**Data S3. (separate file)**

**Overview of genomes used for this study**. Assembly statistics (overall size, number of contigs, N50, estimated completeness) and important genomic characteristics (G+C content, coding density number of CDS) as well as the clade assignment are presented.

**Data S4. (separate file)**

**Results of the ancestral reconstruction using gene tree-species tree reconciliation**. Inferences by ALEml_undated for gene family presence/absence in ancestral and extant lineages, with a general threshold of 0.3 applied.

**Data S5. (separate file)**

**Summary of the ancestral reconstruction**. Inferences by ALEml_undated for ancestral and extant gene family numbers as well as estimated numbers of originations, duplications, transfers and losses per node. Summary of Data S4.

**Data S6. (separate file)**

**Annotation and presence/absence of selected gene families**. Annotation of gene families related to selected biochemical processes are presented, with number of genes as found in the selected Rickettsiales and alphaproteobacterial taxa as well as inferred for the ancestors of Rickettsiales (LRCA), host-associated Rickettsiales (LhRCA) and all Rickettsiales families. Last common ancestor of Anaplasmataceae, Midichloriaceae, Deianiraea and Gamibacteraceae: LMiGDACA; Last common ancestor of Anaplasmataceae and Midichloriaceae: LMiACA; Last common ancestor of Anaplasmataceae: LACA, Midichloriaceae: LMiCA, Gamibacteraceae: LGCA; Rickettsiaceae: LRiCA, Mitibacteraceae: LMCA, Athabascaceae: LAtCA.

**Figshare repository (10.6084/m9.figshare.c.5494977):**

All protein clusters fasta format, proteins alignments, phylogenetic trees generated in the present study

- Vir subunits phylogenetic trees in nexus and pdf format
- Raw ancestral genome reconstruction (ALE) results
- Effector domain profile search (hmmsearch) output
- Protein-level annotations

