## Supplementary material for "The evolutionary origin of host association in an ancient bacterial clade": DataS1: README.txt

Phylogenomics trees README


# RP15_bacteriaBackbone_untr.aln.raxml.treefile
Phylogenetic tree depicting the phylogenetic diversity of MAGs labeled as "Alphaproteobacteria" or "Rickettsiales" in the studies of Parks et al, Tully et al, Delmont et al and Anantharaman et al, as well as the diversity in the contigs of the two re-assembled metagenomes (see Methods). Tree is based on the concatenation of at least 5 out of 15 conserved ribosomal proteins that co-occur on the same contig or MAG. Reference taxa span most bacterial phyla, as well as an archaeal outgroup

# RP15_alphamitoBackbone_f200.aln.iqtree.treefile
Phylogenetic tree depicting the alphaproteobacterial and mitochondrial phylogenetic diversity of sequences that were classified as Alphaproteobacteria in the RP15 tree with bacteria and archaea as reference sequences (see above and Methods). Reference taxa span known alphaproteobacterial diversity as well as mitochondria, Magnetococcales, MarineProteo1 and Beta- and Gammaproteobacterial outgroups. This is the same tree as the one shown in Fig. S1.

# rick116_allTaxa_untreated.aln.pmsftree.treefile.fullnames
Rickettsiales species tree based on the untreated concatenation of 116 highly conserved Rickettsiales aligned marker genes. This tree still includes long branching taxa Candidatus Xenolissoclinum pacificiensis and Candidatus Fokinia solitaria and unstable taxa endosymbiont of Stachyamoeba lipophora and UBA6177. These were removed in further downstream analyses. Tree was inferred under the PMSF approximation of the LG+C60+F+G4 model with IQ-TREE with a 100 non-parametric bootstraps (see Methods).

# rick116_untreated.chain1.con.tre.fullnames
Rickettsiales species tree based on the untreated concatenation of 116 highly conserved Rickettsiales aligned marker genes. This tree is a consensus tree of the first MCMC chain. Tree was inferred under the CAT+GTR+G4 model with Phylobayes-MPI. Consensus tree over all four chains is shown in Fig. S4.

# rick116_untreated.chain2.con.tre.fullnames
Rickettsiales species tree based on the untreated concatenation of 116 highly conserved Rickettsiales aligned marker genes. This tree is a consensus tree of the second MCMC chain. Tree was inferred under the CAT+GTR+G4 model with Phylobayes-MPI. Consensus tree over all four chains is shown in Fig. S4.

# rick116_untreated.chain3.con.tre.fullnames
Rickettsiales species tree based on the untreated concatenation of 116 highly conserved Rickettsiales aligned marker genes. This tree is a consensus tree of the third MCMC chain. Tree was inferred under the CAT+GTR+G4 model with Phylobayes-MPI. Consensus tree over all four chains is shown in Fig. S4.

# rick116_untreated.chain4.con.tre.fullnames
Rickettsiales species tree based on the untreated concatenation of 116 highly conserved Rickettsiales aligned marker genes. This tree is a consensus tree of the fourth MCMC chain. Tree was inferred under the CAT+GTR+G4 model with Phylobayes-MPI. Consensus tree over all four chains is shown in Fig. S4.

# rick116_untreated.pmsfboot.aln.treefile.fullnames
Rickettsiales species tree based on the untreated concatenation of 116 highly conserved Rickettsiales aligned marker genes. Tree was inferred under the PMSF approximation of the LG+C60+F+G4 model with IQ-TREE with a 100 non-parametric bootstraps (see Methods).

# rick116_step47.chain1.con.tre.fullnames
Rickettsiales species tree based on the concatenation of 116 highly conserved Rickettsiales aligned marker genes of which the most heterogeneous sites have been removed. The latter was done by executing 47 rounds of removing the top 1% heterogenous sites according to the chi-square trimmer.  This tree is a consensus tree of the first MCMC chain. Tree was inferred under the CAT+LG+G4 model with Phylobayes-MPI. Consensus tree over all four chains is shown in Fig. S6.

# rick116_step47.chain2.con.tre.fullnames
Rickettsiales species tree based on the concatenation of 116 highly conserved Rickettsiales aligned marker genes of which the most heterogeneous sites have been removed. The latter was done by executing 47 rounds of removing the top 1% heterogenous sites according to the chi-square trimmer.  This tree is a consensus tree of the second MCMC chain. Tree was inferred under the CAT+LG+G4 model with Phylobayes-MPI. Consensus tree over all four chains is shown in Fig. S6.

# rick116_step47.chain3.con.tre.fullnames
Rickettsiales species tree based on the concatenation of 116 highly conserved Rickettsiales aligned marker genes of which the most heterogeneous sites have been removed. The latter was done by executing 47 rounds of removing the top 1% heterogenous sites according to the chi-square trimmer.  This tree is a consensus tree of the third MCMC chain. Tree was inferred under the CAT+LG+G4 model with Phylobayes-MPI. Consensus tree over all four chains is shown in Fig. S6.

# rick116_step47.chain4.con.tre.fullnames
Rickettsiales species tree based on the concatenation of 116 highly conserved Rickettsiales aligned marker genes of which the most heterogeneous sites have been removed. The latter was done by executing 47 rounds of removing the top 1% heterogenous sites according to the chi-square trimmer.  This tree is a consensus tree of the fourth MCMC chain. Tree was inferred under the CAT+LG+G4 model with Phylobayes-MPI. Consensus tree over all four chains is shown in Fig. S6.


# rick116_step47.pmsfboot.treefile.fullnames
Rickettsiales species tree based on the concatenation of 116 highly conserved Rickettsiales aligned marker genes of which the most heterogeneous sites have been removed. The latter was done by executing 47 rounds of removing the top 1% heterogenous sites according to the chi-square trimmer. Tree was inferred under the PMSF approximation of the LG+C60+F+G4 model with IQ-TREE with a 100 non-parametric bootstraps (see Methods).
